# Evaluation and resolution of many challenges of neural spike-sorting: a new sorter

**DOI:** 10.1101/2021.01.19.427297

**Authors:** Nathan J. Hall, David J. Herzfeld, Stephen G. Lisberger

## Abstract

We evaluate existing spike sorters and present a new one that resolves many sorting challenges. The new sorter, called “full binary pursuit” or FBP, comprises multiple steps. First, it thresholds and clusters to identify the waveforms of all unique neurons in the recording. Second, it uses greedy binary pursuit to optimally recognize the spike events in the original voltages. Third, it resolves spike events that are described more accurately as the superposition of spikes from two other neurons. Fourth, it resolves situations where the recorded neurons drift in amplitude or across electrode contacts during a long recording session. Comparison with other sorters on real and simulated ground-truth datasets reveals many of the failure modes of spike sorters. We suggest a set of post-sorting analyses that can improve the veracity of neural recordings by minimizing the intrusion of those failure modes into analysis and interpretation of neural data.

## Introduction

Neuroscientists now routinely apply complex computer algorithms to identify and classify the spikes (“spike sorting”) from multiple neurons in voltage traces recorded by probes that have from one to thousands of electrically-sensitive contacts. The goal of spike sorting is to identify all the spikes that come from neurons near the probe and to cluster those spikes according to the neuron they belong to (Carlson and Carin 2019; Rey et al. 2015). Insofar as the sorting is done correctly, the resulting data allow deep insight into how information is encoded by populations of neurons and also into the connections among the neurons in local micro-circuits. The power of successful identification of the action potentials from many individual neurons simultaneously is indisputable.

Spike-sorters—the algorithms used to perform spike sorting—face many challenges, and it is difficult to compare or rank them. Different sorters exist at different locations in the space of the competing constraints of spike-sorter challenges. Further, a given research goal will require that some challenges of spike-sorting take precedence, while other known failure modes are tolerable. For example, a research project that plans to draw conclusions based on spike-timing cross-correlograms between nearby neurons will need to use a spike sorter that resolves all situations where the waveforms produced by spikes from two neurons overlap (Pillow et al. 2013). Overlaps are a general issue when multiple nearby neurons fire simultaneously either because they are responding to a discrete event, or simply by chance because the neurons have high spontaneous firing rates. Because of our interest in the brainstem and cerebellum, we needed a spike sorter that was ideal for resolving spike overlaps with high reliability for multiple neurons that have firing rates close to 100 spikes/s. We present such a sorter here, called “full binary pursuit” or “FBP”.

We designed FBP to have five characteristics that would allow us to reach our scientific goals. ***1)*** When neurons fire at a high rate and spike overlaps occur with a high probability, the sorter should resolve the overlaps and assign the overlapping spike times to the correct neurons. ***2)*** The sorter should degrade gracefully as signal-to-noise ratio decays to levels too low for reliable sorting. ***3)*** It should be resilient to drift across a long recording session when spike waveforms can change in size or shape and shift gradually from one contact to another. ***4)*** It should avoid creation of false or contaminated output units that are either combinations of real neurons and noise or multi-unit activity from more than one neuron. ***5)*** Failures should be as straightforward as possible to identify and correct during both automated and manual post-processing steps. FBP incorporates these characteristics by borrowing pieces of existing sorters, modifying and combining them, and adding procedures to generate a spike-sorting pipeline that achieves these objectives.

We set out to enable our own scientific goals, but we discovered many cautions about spike-sorting as we validated FBP and compared its strengths and weaknesses with those of three other spike sorters in common use: MountainSort4 (Chung et al. 2017), SpyKING-CIRCUS (Yger et al. 2018), and KiloSort2 (Pachitariu et al. 2016). As a result, our goal here is not to demonstrate that FBP is “better” than the other sorters, but rather to develop a framework to consider the failure modes of spike sorting and strategies for avoiding them. We conclude that “*caveat emptor*” should be the slogan of anyone using a spike-sorter. It is critical to understand the failure modes of a sorter, consider how the failures may or may not compromise the specific scientific goals of a project, and deploy post-processing analyses and criteria that assist in identifying those failures.

## Results

We start with a broad overview of the sorting algorithm, leaving a detailed treatment to the ***Methods*** section. We then assess the performance of FPB by comparison to three other sorters that are in wide use: SpyKING-CIRCUS, MountainSort4, and KiloSort2. Our goal is not to demonstrate that FBP is better than these other sorters (although it is for our needs). Indeed, different sorters make different kinds of errors. Instead, our goal is to highlight the issues faced when using spike sorters and the possible contaminants those issues can introduce into analyses. We consider the challenges of overlaps between the spikes of different neurons, performance under different conditions of signal-to-noise ratio (SNR) in the recordings, drift in space and time, the identification of contaminanted units, and finally various combinations of pitfalls revealed by comparing the performance of all 4 spike sorters on “ground-truth” data.

### Algorithm overview

The FBP sorter has two distinct steps. A “clustering” step thresholds the raw data and goes through a series of computations designed to discover templates across recording channels for each of the neurons in the recording. Then, a “template matching” step uses the templates to find the individual spike event times that correspond to each of the neurons discovered by the clustering algorithm. Spikes are assigned to neurons entirely on the basis of the probabilistic agreement of a snip of the recorded voltages and one of the templates.

#### Clustering

Our sorting pipeline begins by whitening the noise across channels via zero phase component analysis (ZCA). To avoid the problems of drift across space and time, we divide the whitened voltage into shorter time segments, and initially analyze each channel and time segment independently. We threshold the whitened voltage traces to obtain brief spike event clips that form the basis for clustering. We perform principle component analysis (PCA) on the spike clips for each channel and time segment and use an automated reconstruction accuracy based procedure (Abdi and Williams 2010) to select the optimal principle component features. We cluster the spike clips in PCA space, following a modified version of the MountainSort “isocut” clustering algorithm (Chung et al. 2017). Briefly, we massively over-cluster the data and compare each cluster with its mutually nearest neighbor, making a binary statistical decision whether or not to merge the two clusters into a single cluster, or optimally split them into two distinct clusters. This iterative process results in a set of clusters, each corresponding to a set of spike events produced from a putative single neuron during a given time segment on a particular channel.

Next, we combine clusters across channels to obtain a single set of neuron waveform templates for each time segment. We compute the average across the spike clips for each cluster along with the averages across all other channels for the times of the spike clips. Then, we concatenate the averages across channels to represent each cluster as a single template over all channels. Because we discovered the clusters, and thus the templates, independently on each channel, a single neuron might be represented by more than one cluster. We therefore condense the templates by finding the pairs of templates that are most similar to one another and using the same merge test as in the clustering algorithm to ask whether these two clusters should be combined. We repeat the process pairwise until no clusters combine, leaving templates that correspond to the waveforms across channels of a unique set of neurons irrespective of the channel they were detected on.

#### Template matching

The goal of our template matching algorithm is to find the individual spike event times that correspond to each of the neurons discovered by the clustering algorithm. We discard the individual clips used during the clustering step and use the unique templates from the clustering algorithm to perform a slightly modified version of binary pursuit template matching (Pillow et al. 2013). Thus, the spike times output by FBP are determined fully by binary pursuit and not by threshold crossings.

Binary pursuit is a greedy, iterative approach to template matching. Each iteration detects the optimal fit of a template to a site in the voltage, labels that time as a spike, and subtracts the template from the voltage trace to create a residual voltage trace. Template matching has 3 major benefits over thresholding and clustering. (1) Binary pursuit considers information in the recorded voltage across multiple channels and multiple time points to determine the time of a spike and the identity of the neuron that produced it. (2) Multiple spikes overlapping in time can be assigned to multiple neurons, improving detection of overlapping spike events. (3) We include a noise term that equips the algorithm to reject voltage deviations that are more likely to have been generated by noise or artifacts. FBP also includes an explicit procedure to identify and resolve voltage deflections that result from the temporal overlap of spikes from 2 neurons. For each spike event detected by binary pursuit, we ask whether the spike event is better explained as a sum of two templates than as a waveform from the originally assigned template alone.

Last, we stitch the sorted neurons back together across the individual time segments on which the data were sorted. We again use the merge test from our clustering algorithm in pairwise fashion, taking advantage of abundant overlap between adjacent time segments. We also assess the possibility that what appear to be two neurons are actually a single neuron whose recording has drifted across contacts on a multi-contact probe or has changed amplitude across the duration of the recording. Post-sort processing removes any units with excessive refractory period violations in their autocorrelograms, assesses combined autocorrelograms of pairs of units to identify single neurons that have been sorted into two units, and asks whether a set of spikes might have been assigned to two neurons by analyzing crosscorrelograms (CCGs) between pairs of units. If a pair of units exhibits an excessive peak at time zero in their CCG, the unit with lower quality is removed. This renders a final output that minimizes the number of “neurons” that are actually multi-unit activity or mixtures of well-isolated units.

### Resolution of overlapping spikes

Overlapping spikes result from the voltage deviation of two distinct neurons, within roughly a spike width of each other and roughly in the same spatial location across electrodes. They present a problem that has a potential impact on neural data quality and interpretation, especially when spike rates are high. Overlaps can cause spikes to be missed, reduce the recorded firing rate, alter measured responses to a given stimulus, and contaminate neural cross-correlograms. Even for experiments that simply investigate stimulus response properties, nearby neurons frequently share tuning properties and will thus disproportionately tend to fire spikes at the same time. Without resolution of overlaps, spike detection during sorting will be worst during the epoch of greatest experimental interest.

#### Random overlaps

We first consider a simulation with two model neurons that fire independently at rates that roughly mimics our own use case in the cerebellum: one at 60 Hz and the other at 90 Hz. We ran the simulation for 200 s, generating 12,095 and 18,055 spikes respectively (Figure 1a). Both units have Poisson random firing rates with a 1.5 ms refractory period. The data were generated using two templates with largely non-overlapping spatial distributions and very high SNR (10.5 and 6.1; Figure 1c. The two spikes could be sorted perfectly by choosing a threshold beyond the noise envelope and assigning any threshold crossings on channel 1 to unit 2, and any threshold crossings on channel 2 to unit 1 (Figure 1b).

**Figure 1.**
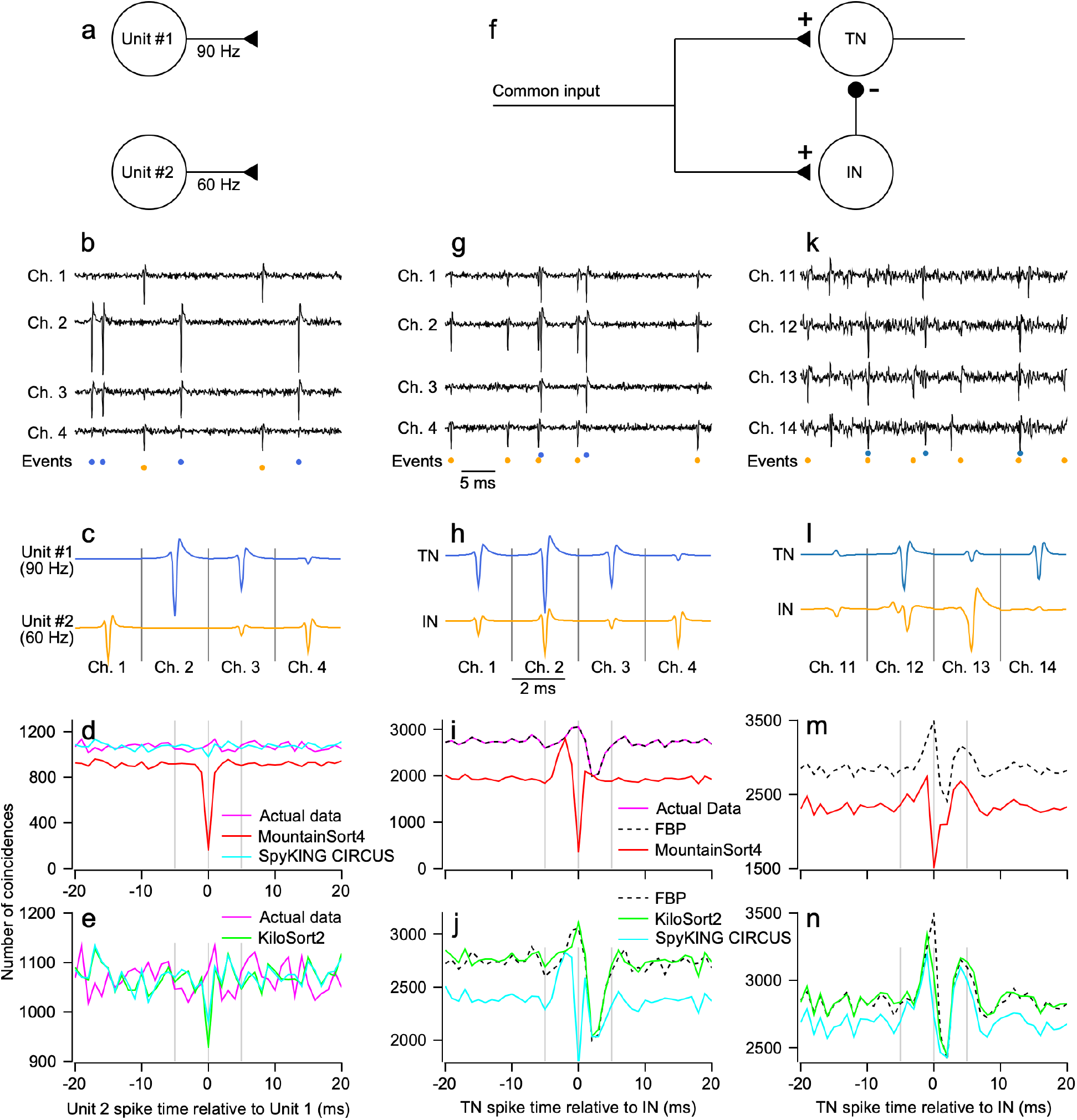
Sorting accuracy for overlapping spikes in synthetic and real data. ***a)*** Simulated network comprised of two, independent units with Poisson firing rates. ***b)*** Sample voltage trace from synthetic tetrode dataset generated from the templates in ***c***. Colored symbols below the traces indicate time points where corresponding templates in ***c*** were added. ***d-e)*** CCGs between the best matching units for each sorter and actual data. FBP output was identical to the actual data and is not shown. ***f)*** Simulated network with two units embedded in a simple feedforward inhibition circuit. ***g-j)*** same as ***b-e*** with spike times taken from the output of the feedforward leaky-integrate-and-fire network. ***k)*** Sample voltage traces from 4 channels of a real recording obtained from the cerebellar cortex using a low-density probe. Colored symbols below the traces show the event times discovered by FBP. ***l)*** Average templates of two units discovered by FBP. ***m-n)*** CCG of the two FBP output units and the two closest matching units from the other sorters.

FBP detected all spikes correctly and produced a CCG that matched the actual dataset exactly (not shown). In contrast, the other 3 spike sorters missed overlapping spikes to some degree (Figure 2d, e). The actual dataset (magenta trace) contained 1085 overlapping spikes at t=0, approximately 9% of unit 1 spikes and 6% of unit 2. MountainSort4 has a powerful clustering algorithm but does not implement a templated matching procedure and produced a CCG with a dip at time zero (Figure 1d, red curve), indicating that it had missed or misclassified almost all overlaps. Both SpyKING-CIRCUS and KiloSort2 take a very similar approach as FBP by performing clustering followed by template matching, and both performed reasonably well. Both detected the majority of overlapping spikes in the simulated dataset (Figure 1d, cyan curve; Figure 1e, green curve). However, a higher gain on the y-axis of the CCG (Figure 1e) reveals that each of these sorters missed in excess of 100 overlapping spikes, nearly 10% of the total overlaps in the dataset. These failures did not impact the overall firing rates greatly but could have altered the shape and size of small responses to a given condition.

**Figure 2.**
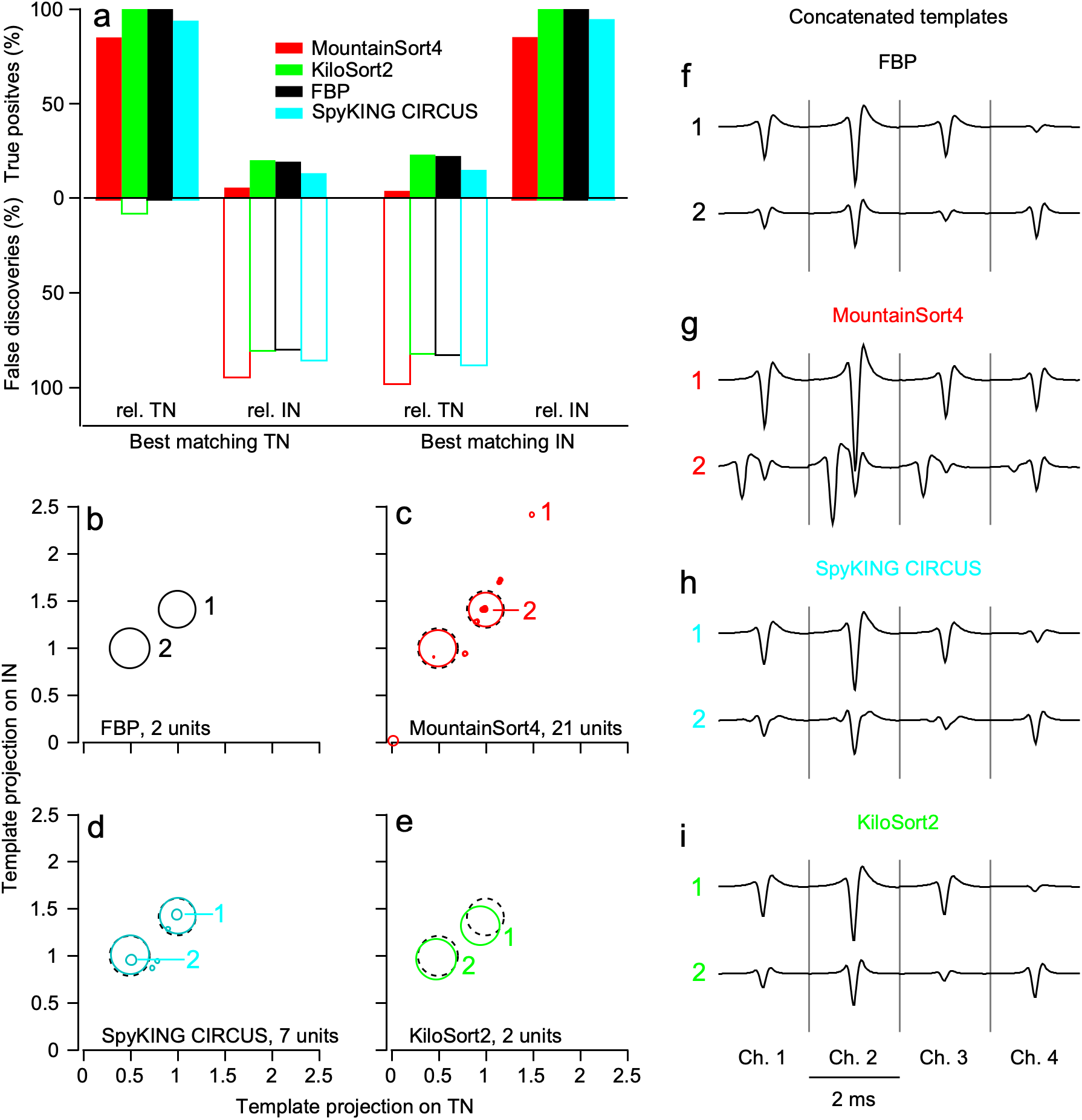
Quantitative analysis of sorting accuracy for synthetic feedforward-inhibition dataset. ***a)*** Histogram bars above and below the abscissa show TP rates and FD rates for the same dataset in Figure 1 *f-j*. Different colors indicate accuracy for different sorters. The left and right pairs of bars show values for the unit best matching the true target neuron (TN) and the true inhibitory neuron (IN), respectively. Within each pair of bars, TP and FD rates are shown for each best matching unit computed against the real TN spike event times and the real IN spike event times. ***b-e)*** Scatterplots of the projection of each unit’s template against the true TN and IN templates for each sorter. The area within each symbol is scaled linearly as a function of the number of spikes discovered in each unit. Dashed circles indicate the projections of the true TN and IN templates, which overlapped perfectly with the projections for FBP and therefore do not appear in panel ***b. f-i)*** Templates for 2 example units discovered by each sorter as labeled in ***b-e***.

#### Performance on a modeled feedforward inhibition motif

We next simulated a feedforward inhibition neural circuit because it comprises a common scenario in physiological recordings. The circuit consists of two leaky integrate-and-fire neurons (see ***Methods***) that receive a powerful shared input of constant excitation with a square wave modulation (Figure 1f). The target neuron (TN) is inhibited directly by the inhibitory neuron (IN). We added our selected neuron templates (Figure 1h) to noise at spike times determined by the model to create the synthetic test data set (Figure 1g). We again used high SNR (TN, 10.5 and IN, 6.1) and very high firing rates (TN, 21,622 spikes, ~108 Hz; IN, 25,155 spikes, ~128 Hz) to remove signal detection confounds and emulate the firing rates of cerebellar neurons or neurons under stimulus drive. Now, the spike times are no longer independent and the neurons’ templates have considerable spatial overlap across channels.

FBP performed perfectly on the simulated data set. It revealed a CCG that matched the actual data (Figure 1i, magenta and black dashed traces). The CCG exhibits the hallmark features of feedforward inhibition with an increase in spike synchrony for several ms before the IN fires followed by inhibition of the TN. By contrast, because MountainSort4 does not consider overlapping spikes, its CCG is severely distorted and the effect of inhibition on the TN is not evident (Figure 1i, red trace). SpyKING-CIRCUS detects many more of the overlapping spikes, yet it produces a CCG that distorts the overall feedforward inhibition motif (Figure 1j, cyan trace). The CCG suggests a much larger proportion of TN spikes preceding an IN spike and a much lower level of inhibition than exists in the actual data. Finally, KiloSort2 largely reproduces the correct CCG (Figure 1i, green trace).

#### Performance on cerebellum recording

We next compared the 4 spike sorters on 10 minutes of actual data recorded from the primate cerebellum using low-density 32 contact linear array electrodes (Figure 1k). The templates from FBP reveal 2 units with a moderate degree of spatial overlap and sufficient SNR for an accurate sort (Figure 1l). The putative TN had a firing rate of 48 Hz and a SNR of 3.5; the putative IN had a firing rate of 94 Hz and a SNR of 4.3. Based on the sort delivered by FBP, the CCGs of these two units exhibit the signature increase, then decrease, of feedforward inhibition circuitry (Figure 1m, dashed black trace). As before, MountainSort4 largely misses overlapping spikes and has a CCG with a dip at t=0 (Figure 1m, red trace). KiloSort2 and SpyKING-CIRCUS (Figure 1n, green and cyan traces) both provide a CCG that is consistent with the CCG from FBP, but with a lower estimate of the number of overlapping spikes. We do not know ground truth in the cerebellum dataset, but the consensus among FBP, SpyKING CIRCUS and KiloSort2 is that the recording contains two units whose firing rates are correlated at fine timescales.

#### True positives and false discoveries

We can learn more about the strengths and weaknesses of a spike sorter by assessing performance in synthetic datasets where spike timing and neuron identity are known. For the model TN and IN used in Figures 1g-j, we computed true positive (TP) rate for each sorted unit relative to each ground truth neuron as:

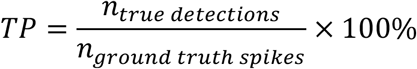

where a sorted unit was credited with a “true detection” if it contained at least 1 spike event time within +/- 1 ms of a ground truth spike time. By this definition, a unit’s TP rate cannot exceed 100%. For example, a unit containing a spike 0.5 ms before *and* after every ground truth spike would contain twice the number of ground truth spikes, but still have a TP rate = 100%. This definition of TP also implies that non-zero TP rates for sorted units that do not match the ground truth neuron are not necessarily errors. Accurately sorted non-ground truth spikes can occur within +/- 1 ms of a ground truth spike and be credited as “true detections”. On the other hand, a TP rate below 100% indicates that ground truth spikes were undetected. Throughout the paper, we match the sorted unit with the highest TP rate to the ground-truth neuron for computing sorter accuracy.

We compliment the TP rate with the false discovery (FD) rate for each sorted unit. We define the FD rate as:

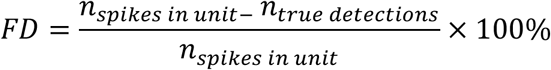

Intuitively, FD rate indicates the percentage of each unit’s spikes that are *not* ground truth spikes: it is the percentage of a unit’s spikes that do not lie within +/-1 ms of a ground-truth spike. The FD rate is necessary because our formula for TPs does not penalize a spike sorter for being overly permissive. In our simplistic example above, the unit containing 2 spikes for every ground truth spike will have a FD rate = 50%, in contrast to its TP rate of 100%. Taken together, an ideal sorter would deliver TP = 100% and FD = 0% for its best matching ground truth unit, and low rates of TPs accompanied by high FD rates for the other units. Again, FD rates under 100% are acceptable because of the way “true detections” are attributed to each unit.

We analyzed the TP and FD rates of the sorters relative to both the IN and TN units that comprise the synthetic feedforward dataset of Figure 1g-j. We define the best matches as the sorted units with the largest TP rates and we call them the “best matching IN unit” and the “best matching TN unit”. The performance of FBP on this dataset was perfect and can be taken as the ground truth sorting result (Figure 2a, black bars). KiloSort2 (Figure 2a, green bars) has a TP rate of nearly 100% for both its best matching TN and IN units. The TP rates were ~10-15% for both FPB and KiloSort2 for the best matching IN unit relative to the true TN and the best matching TN unit relative to the true IN, as expected given the actual number of overlapping spikes. MountainSort4 (Figure 2a, red bars) and SpyKING CIRCUS (cyan bars) both had TP rates below 100% for the best matching TN unit and the best matching IN unit, confirming the poor detection of overlapping spikes indicated by the CCG analysis.

FD rate also is also critical to understanding sorter performance. Only KiloSort2 has a non-zero FD rate (7.3%) for the best matching TN (Figure 2a, open green bar). Because KiloSort2 had 99.9% TPs for both the TN unit and the IN unit, nearly all ground truth spikes are accounted for. The presence of FDs therefore indicates an excess of spikes added to the best matchingTN unit, rather than incorrect assignment of true IN spikes to the TN unit. For the other sorters, the TN and IN units were generally free of FDs (leftmost and rightmost sets of open bars). Finally, as expected, all sorters have FD rates close to 100% for the best matching IN unit relative to the true TN and vice versa.

#### Sorted templates versus ground-truth

We further quantified how well the sorters classified ground truth spikes by comparing the average template for each unit identified by each sorter with the ground truth templates. This step is necessary because the TP rates alone for a sorted unit do not reveal the extent to which TP detections are due to a spike sorter correctly detecting a ground truth spike as opposed to another event occurring nearby. We reasoned that if a sorted unit’s spike events are the direct result of the detection of ground truth spikes, then voltage clips centered on each spike time should be consistently aligned with ground truth spikes. Thus, the average voltage trace centered on each spike time (i.e. the template) should look like the ground truth template to the extent that a unit’s TP detections directly result from the detection of ground truth spikes.

The four scatterplots in Figure 2b-e provide a succinct visual summary of how well the templates found by the 4 spike sorters matched the ground-truth templates. We quantified the similarity between each unit’s template and the ground truth using the scalar projection between templates. The scalar projection was taken as the peak value of the cross correlation function between the templates to allow for small alignment differences between sorters. The scalar projections then were normalized by the magnitude of the ground truth template so that the true TN and IN lie at (1, y) and (x, 1). Each point in the scatterplots shows the template projection onto the TN and IN templates for one unit. The area of each circle is linearly related to the number of spikes placed in each unit. FBP (Figure 2b), MountainSort4 (Figure 2c), and SpyKING-CIRCUS (Figure 2d) all found two units that overlay the ground truth template projections (black dashed circles) almost perfectly, indicating that these units represent correct classification of ground truth spikes. The template for the best matching TN unit for KiloSort2 is shifted from the ground truth TN (Figure 2e), in a direction that suggests it looks more similar to the IN than expected.

Direct inspection of the templates suggests the types of errors made by the sorters (Figure 2f-i). The templates for KiloSort2 (Figure 2i) are very similar to the ground truth templates (Figure 2f) except that the TN unit template has a slightly diminished amplitude. Together with the high FD rate for the TN, the shifts in template amplitude and location suggest that KiloSort2 double counted some of the IN spikes and classified them as TN spikes. Both MountainSort4 and SpyKING-CIRCUS output more than 2 units (21 and 7, respectively), likely because they mishandled overlapping spikes and placed them into separate clusters. For MountainSort4 (Figure 2g), example template 1 results from nearly directly overlapping TN and IN spikes while example template 2 shows overlapping spikes at a longer time lag. For SpyKING-CIRCUS (Figure 2h), the two templates shown are close to the correct location in the scatterplot. However, there are extra units with a small number of spikes whose templates lack the smoothness expected from the ground truth template, probably because of poor spike timing alignment and confusion from overlapping spike events.

### Sorting under reduced SNR

All spike-sorters will eventually fail as SNR decreases toward zero. One goal of FBP is to remain roughly as likely to miss non-overlapping and overlapping spikes across a range of SNR. Here, we evaluate the performance of FBP and KiloSort2 on 3 synthetic datasets that vary in SNR but otherwise contain the same spike times from the TN and IN neurons from our feedforward inhibition model. We chose templates that are more difficult to distinguish from one another (Figure 3a-c) and varied the SNR of the templates across 3 levels (SNR = 8.7, 4.4, 2.6 for the TN and 6.6, 3.3, 2.0 for the IN). SNR is taken from the channel with maximal SNR according to equation 8.

**Figure 3.**
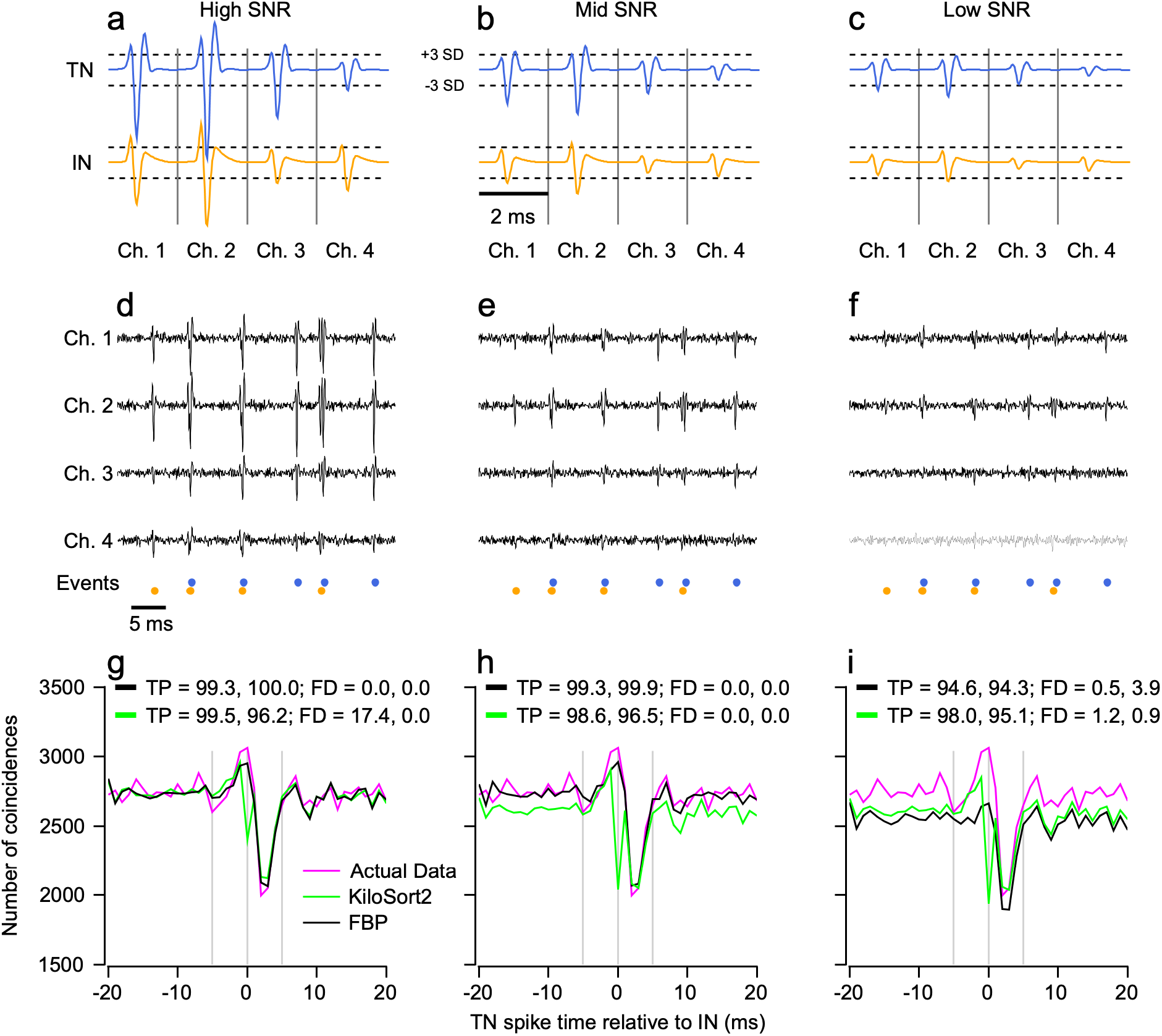
Detection of overlapping spikes and sorter performance as signal-to-noise ratio degrades. ***a-c)*** Templates used to generate the synthetic spike trains. Dashed horizontal lines indicate 3 standard deviations of the noise used to create the synthetic data. ***d-f)*** Example voltage traces for each SNR. Spike event times are the same as in the synthetic feedforward inhibition dataset of Figure 1 ***f-j. g-i)*** CCGs constructed by plotting the distribution of the spike times for the best matching TN unit vs. relative to the time of the best matching IN spike for KiloSort2 and FBP for each level of SNR. Values of TP and FD given in each graph are for the TN and IN, in that order.

FBP performs extremely well under conditions of high SNR (Figure 3, left column). It detects 100% of IN spikes and 99.3% of TN spikes without any FDs for either unit. The sorted units reproduce the ground truth CCG well (Figure 3g, black versus magenta traces), including the peak at time 0. KiloSort2 misses about 4% of IN spikes and detects 99.5% of TN spikes. The CCG shows a large dip at time zero (Figure 3g, green curve) suggesting that overlapping spike events are most likely to be missed.

Performance is similar for the middle SNR (Figure 3, middle column): FBP misses a small percentage of spikes, has zero FDs, and reproduces the ground truth CCG fairly accurately (Figure 3h, black versus magenta curves). KiloSort2’s TP rates are very similar to those for a high SNR and, interestingly, it no longer has a large number of FDs. KiloSort2’s bias toward missing overlapping spikes has increased further, causing a larger dip in the CCG at time zero (Figure 3h, green curve).

The performance of both sorters degrades for low SNR (Figure 3, right column). TP rates fall and FD rates increase, although the errors are generally under 6%. Here, FBP detects fewer spikes overall than does KiloSort2. However, the CCGs (Figure 3i) indicate that FBP’s detection of overlapping spikes is not degraded selectively as there is not a large dip in the CCG at time 0. In contrast, KiloSort2 has become further biased and its dip at time 0 has increased in magnitude. Overall, the performance of FBP in detecting overlapping spikes is stable in comparison to its detection of non-overlapping spikes, despite the fact that it detects fewer overall spikes than KiloSort2 in low SNR conditions.

### Compensation for drift in space and time

In actual neural recordings, the waveforms of individual neurons may drift in amplitude and/or move to different channels across time. FBP compensates for drift by explicitly dividing the recording into short adjacent segments that overlap in time. Sorted units in one segment are joined with units in an adjacent segment if their spike clips merge together via our clustering algorithm or their spike times are largely identical within the interval of overlap. We chose to perform drift compensation by working in time because it is a directly measured variable during neural recordings and the investigator has a good intuition for adjusting the length of segments and their degree of overlap. Here we benchmark our drift correction procedure against that of KiloSort2, which is the only other sorted we tested with an explicit drift correction procedure that is conducted in PCA space.

To evaluate compensation for spatial and temporal drift, we generated a synthetic dataset with two ground truth units defined on 8 voltage channels. A “static” neuron fired spikes at 60 Hz and did not drift (Figure 4a, b, orange templates) and a “drift” neuron fired spikes at 90 Hz and had a template whose amplitude across channels drifted rapidly over time (Figure 4a,b, magenta templates); the speed and magnitude of the drift exaggerate what happens in real life, but simplify our analysis and allow us to work with only 120 s of data. Both units had independent Poisson random spike times with a 1.5 ms refractory period. The drift neuron’s template started as shown in Figure 4a at time t = 0, with a peak amplitude on channel 5 and ended at time t = 120 s with a peak amplitude on channel 6.

**Figure 4.**
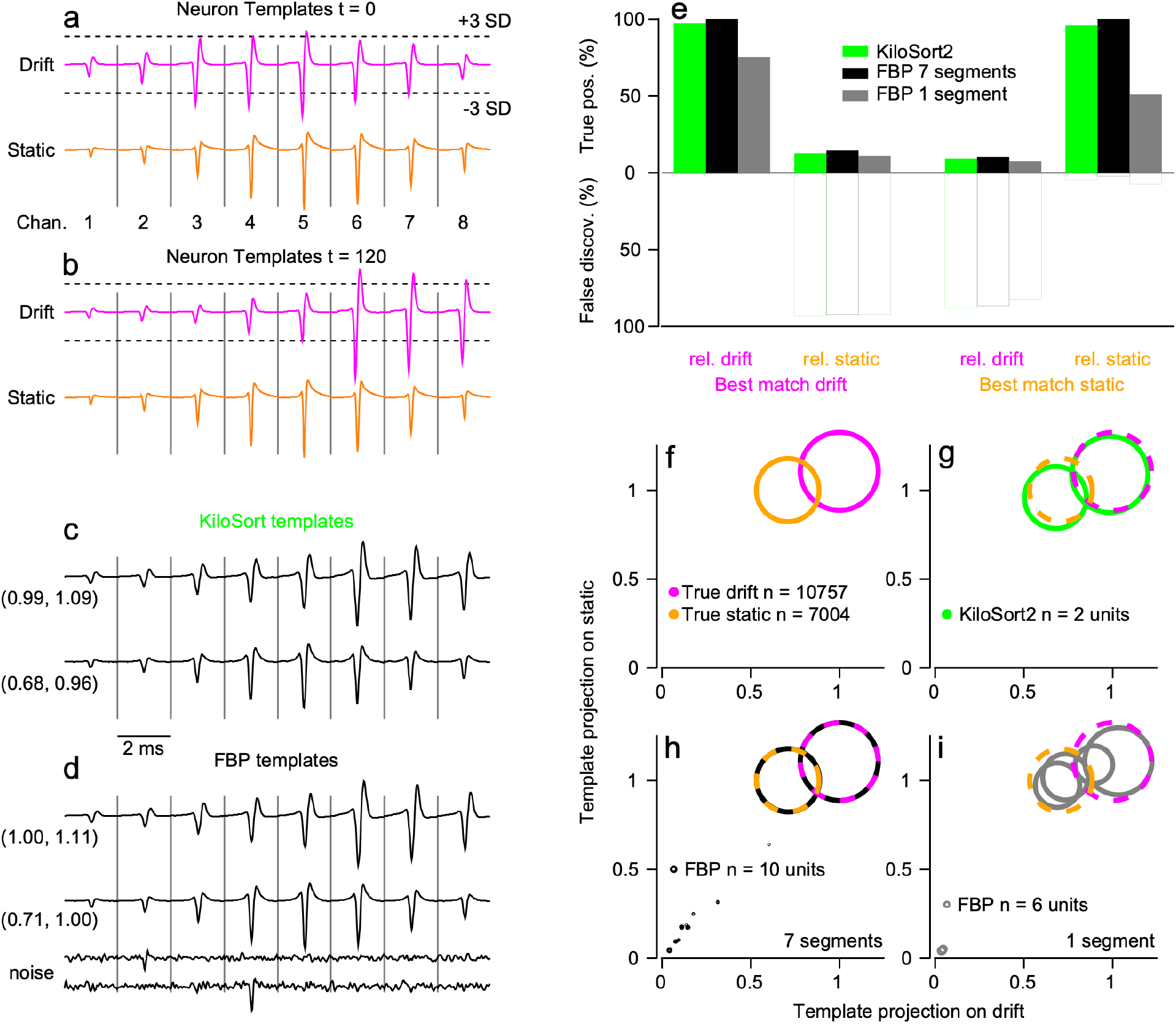
Drift correction for a synthetic dataset. ***a)*** Unit templates used to add spikes at the beginning (time = 0 s) of the synthetic dataset for the drifting unit and static unit. ***b)*** Unit templates used to add spikes at the end of the dataset (time = 120 s). The drift template has moved across channels and the static template has remained stable and therefore is the same as in ***a. c-d)*** Templates of the units output by KiloSort2 using its PC-based drift correction and and by FBP using its time-based drift correction by dividing the data into 7 overlapping segments. Coordinates to the left of each template indicate their location in scatterplots in ***f-i***. FBP noise examples are in the lower left of ***d. e)*** TP and FD rates for each sorter’s best matching drift and static units. Conventions are the same as Figure 2a. Black bars show FBP results using 7 time-segments for drift correction while gray bars show FBP results for 1 time segment, that is without drift correction. ***f-i)*** Scatterplots of the projection of each unit’s template against the true static and drift templates for each sorter. The area within each symbol is linearly related to the number of spikes in its corresponding unit. Dashed circles in ***g-i*** indicate the projections of the true static and drift templates for reference.

KiloSort2 performs well even under rapid drift rates and with interference from the static unit (Figure 4e, green bars; drift unit: 96.0% TP, 0.0% FD; static unit: 94.8% TP, 3.6% FD). FBP performs almost perfectly with a segment duration of 30 seconds to account for the rapid time scale of the drift and 15 s overlap between segments (Figure 4e, black bars). By contrast, FBP performs poorly when sorting the drifting dataset with a single time segment, disabling explicit compensation for drift (Figure 4e, gray bars).

Analysis of the templates produced by averaging the spikes detected by each sorter tells a similar story (Figure 4c,d). The best-matching static and drift templates for both KiloSort2 and FBP are qualitatively very similar to the true templates. The quantitative template projections show that the KiloSort2 templates are not perfect matches for the ground-truth templates (Figure 4g), confirming a small degree of sorting or alignment errors. The templates for FBP project perfectly onto the ground-truth templates with drift compensation enabled (Figure 4h) but poorly when data were sorted in a single segment (Figure 4i). Without drift compensation, FBP splits the drifting and static neurons into multiple units with smaller numbers of spikes in each unit (Figure 4i, gray circles). Thus, both sorters perform well in the face of severe spatial-temporal drift. Our approach working in the time domain can match or exceed KiloSort2’s approach in PC space, at the cost of being more computationally expensive.

We note that FBP finds 8 extra units even with drift compensation enabled. These units contain a very small number of spikes and are identified easily as the result of noise deflections in the dataset. Examples of the noise unit templates appear at the bottom of Figure 4d and their templates do not project near ground truth (Figure 4h, small black circles).

### Parsing true versus false units in real neural recordings with ground truth

Typical neural recordings from multi-contact electrode arrays yield complex datasets that pose substantial challenges to spike sorting algorithms and the assessment of sorting quality. In real neural recordings, many neurons are present across multiple contacts, and their isolation quality, SNR, and similarity to nearby neurons spans a spectrum of possibilities. Spike sorter confusion between individual neurons and noise is inevitable, leading all sorters (including FBP) to the discovery of “false units” that do not represent any neurons actually present in the data. We developed FBP with a bias to output units that are well separated from each other and make fewer errors of confusion, at the expense of being more likely to split true neurons or miss spikes. Our assumption was that we could not exclude false units entirely, so we targeted a strategy that would allow us to most easily identify false units due to multiunit activity or contamination by noise in automated and manual post-processing steps.

We explored the identification of true and false units by the 4 spike sorters on the “ground-truth” “PAIRED_KAMPFF” dataset contributed to SpikeForest2 (Magland et al. 2020) by the Adam Kampff lab (Marques-Smith et al. 2018; Neto et al. 2016). This dataset consists of a patch clamp recording from one neuron aligned with simultaneous extracellular recordings using a dense Neuropixels microelectrode array. The extracellular data comprised 32 channels surrounding the channel with the best extracellular isolation of the intracellularly recorded ground truth unit. This dataset poses both an opportunity and a challenge. The opportunity is to determine how well each sorter can identify the ground-truth neurons and to understand their failings. The challenge is that the recordings include spikes from many neurons and the sorters find many units. Even if sorting accuracy is excellent for the ground-truth neuron, there is no way to assess sorting accuracy for the other units found by the sorters. Focusing solely on the accuracy for the ground-truth unit risks overestimating overall sorting quality.

We take a more holistic approach and attempt to gain insight into the separation between the ground truth unit and other sorted units by evaluating sorting accuracy across all units discovered by a sorter. Our evaluation begins by considering 5 ground truth datasets and identifying the top 10 units discovered by the sorters in decreasing order of TP detections of the ground truth spikes (Figure 5). If we consider only the unit with the highest TP rate (unit 0), then with a couple of exceptions, the output units for all sorters and datasets in Figure 5 look good to excellent, with TP rates near 100% and FD rates near 0%. However, inspection of the TP rates across the next 9 units reveals that in many cases, non-ground truth units have fairly high TP rates. An ideal spike sorter will yield a binary result: output units are either the ground truth neuron or not. The best matching unit (unit 0) should have 100% TPs and 0% FDs and the remaining units 1-9 should have low TPs and high FDs. Even in a perfect sort, non-ground-truth units will exhibit non-zero TPs due to temporal coincidence of their spikes within +/- 1 ms of ground truth spikes. However, coincident spikes near a rate of 25% seems physiologically implausible: it implies an extraordinary level of neuron-neuron spike correlation with a very high level of temporal precision. High TP rates and/or low FD rates in units beyond the ground-truth unit 0 suggest the presence of false units.

**Figure 5.**
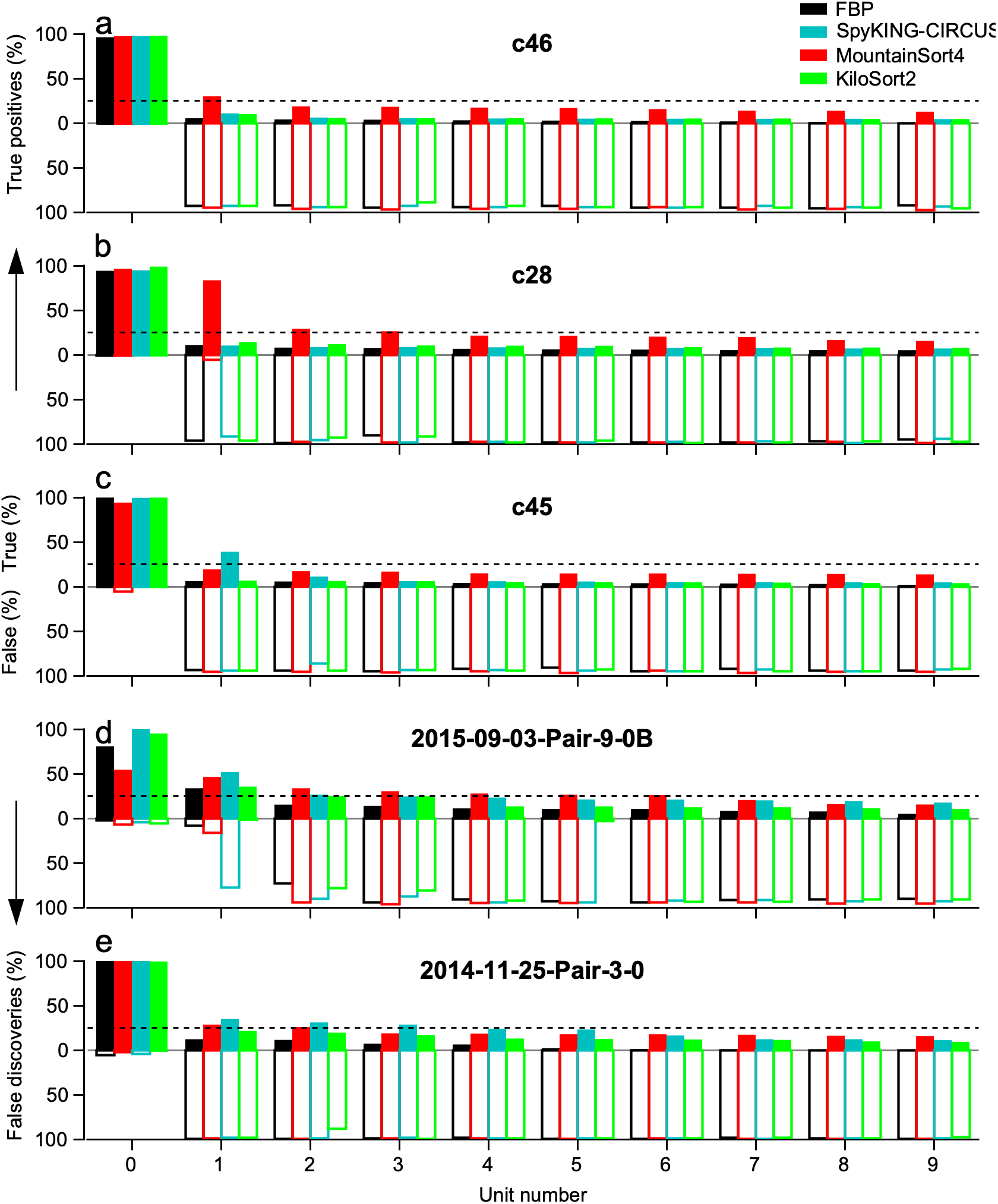
Quantitative assessment of spike sorter accuracy for 5 files of the SpikeForest Kampff dataset. Each graph presents data for a different recording and plots the TP and FD rates ordered by TP rate for the 10 units with highest TP detections of the ground truth unit. Bars above the abscissa (filled) show TP rates, those below the abscissa (unfilled) show FD rates. Data are shown for each of the 4 sorters tested. FBP, black; SpyKING-CIRCUS, cyan; MountainSort4, red; KiloSort2, green. The SNR and maximum SNR channel (see METHODS) for each recording are: file c46, 5.25 on channel 17; file c28, 6.70 on channel 16; file c45, 5.65 on channel 16; file 2015-09-03-Pair-9-0B, 12.50 on channel 9; file 2014-11-25-Pair-3-0, 10.50 on channel 3.

Consider, for example, the performance of MountainSort4 for file c28 (Figure 5b). Unit 0 is an excellent representation of the ground-truth neuron. But the high TP and low FD rates of unit 1 indicate it is a redundant copy of the ground truth neuron. Units 2-9 have TP rates larger than those of the units discovered by every other sorter coupled with high FD rates. This pattern suggests that many units discovered by MountainSort4 have an excess of spikes that overlap the ground truth spikes, suggesting that these units are either MUA that includes ground-truth spikes and/or other units contaminated by noise. For the dataset 2014-11-25-Pair-3-0 (Figure 5e), SpyKING-CIRCUS has a near perfect sort of the ground truth neuron but shows a similar pattern of high TPs and FDs for units 1-6. In the same dataset KiloSort2 unit 2 has over 19% TPs and only 88% FD, suggesting a unit that either is very strongly correlated with the ground truth unit or is a mixture of ground truth spikes with other events. The preponderance of high values of TP rate for units 1-9 across the datasets in Figure 5 suggests the presence of many false units. FBP has the lowest TP rates among units 1-9 almost uniformly. We suggest that the lack of TPs in the sorts from FBP reflects fewer false positive units and good separation between the ground truth unit and other neurons in the recording. It is unlikely to be the result of poor detection of coincident spikes because FBP outperforms the other sorters in resolving overlapping spikes (Figure 1).

We dig deeper into the issue of false units using a dataset that gave all 4 sorters difficulty, file 2015-09-03-Pair-9-0B from Kampff dataset (Figure 6). FBP appears to sort the ground truth unit poorly (Figure 6a). Unit 0 has a TP rate of 54% and a FD rate of 7%: it is missing a large number of ground truth spikes and it contains spikes that do not correspond to the ground truth neuron. Unit 1 has a TP rate of 46% and 1% FDs: 99% of its spikes correspond to the ground truth neuron but a large number of spikes are missing. The remaining FBP units, 2-9 show much lower TP rates, indeed the lowest of all 4 sorters and very high FD rates. Together, the TP and FD rates suggest that FBP split the ground truth unit in half in its output, but otherwise successfully isolated the ground truth unit.

**Figure 6.**
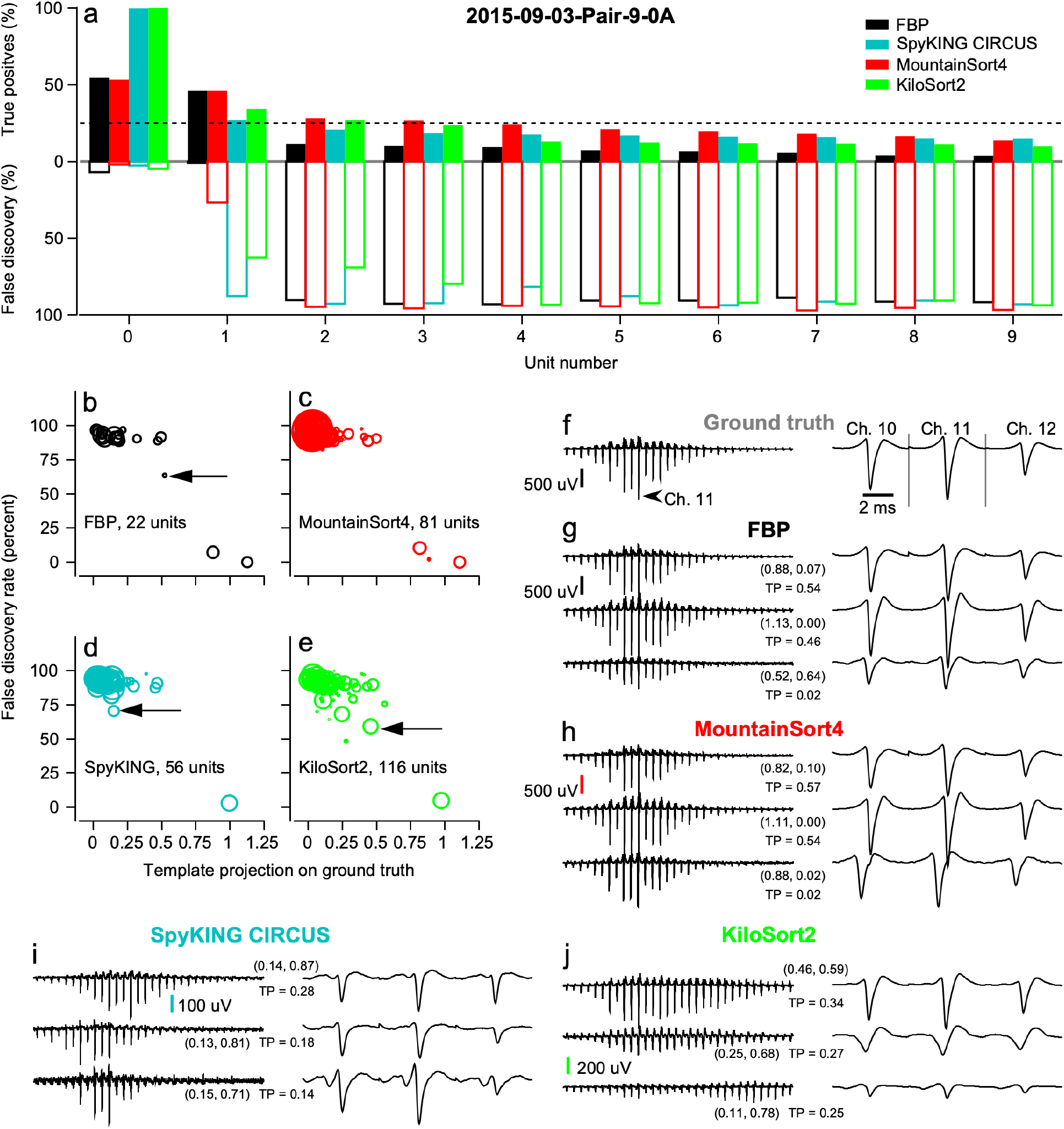
Detailed assessment of spike sorter performance on ground truth data from the SpikeForest Kampff 2015-09-03-Pair-9-0A dataset. ***a)*** The graph plots the TP and FD rates ordered by TP rate for the 10 units with highest TP detections of the ground truth unit. Bars above the abscissa (filled) show TP rates, those below the abscissa (unfilled) show FD rates. Data are shown for each of the 4 sorters tested. FBP, black; SpyKING-CIRCUS, cyan; MountainSort4, red; KiloSort2, green. ***b-e)*** Scatterplots of FD rate vs. normalized template projection on the ground truth template for all units output by each sorter. The actual ground truth unit (not shown) would plot at (0, 1). Arrows point to symbols for the units that show evidence of greatest spike sorter error. The area within each symbol is linearly related to the number of spikes discovered for its corresponding unit. ***f-j)*** Three example templates from each sorter. In the left column, the templates include all 32 channels in the dataset on a slow time scale. In the right column, the templates show the same data on a faster time scale for 3 channels centered on the channel of the largest spike from the ground truth unit. Coordinates next to each template indicate their location in scatterplots in ***b-e***. The ground truth SNR is 10.00 on channel 11.

Performance for the other three sorters is varied. For the most part, the TP rates of units 1-3 are higher than we would expect from random overlap with the ground-truth spikes (Figure 6a). MountainSort4 (Figure 6a, red bars) appears to split the ground truth neuron in a manner similar to FBP. However, its unit 1 has a high 27% FD rate suggesting that the unit is contaminated by non-ground truth spikes. Units 2 and 3 have TP rates in excess of 25% (dashed horizontal line) and very high FD rates, suggesting that they represent multi-unit activity due to a large number of ground truth spikes plus many other events. SpyKING-CIRCUS’s (Figure 6a, cyan bars) unit 0 is a near perfect representation of the ground truth neuron with nearly 100% TPs and 0% FDs. However, units 1 and 2 contain 27% and 21% ground truth spikes and have high FD rates implying that these 2 units are likely to be mixtures that include a large number of ground truth spikes. KiloSort2’s (Figure 6a, green bars) unit 0 is also a near perfect representation of the ground truth neuron. However, units 1-3 have TP rates of 34%, 27% and 24% and relatively low FD rates, implying that the ground truth spikes comprise a large proportion of their spike events.

To understand the situation better, we again computed the ground truth template (Figure 6f) and the average concatenated templates for all units. We projected the template for each unit onto the template of the ground truth unit and made scatterplots that are slightly different from the ones we used in Figures 2 and 4 (Figure 6b-e). Here, we plotted the FD rate versus the template projection on ground truth. In these scatterplots, an ideal sorter would deliver one unit with a FD rate of 0% and a template projection of 1, corresponding exactly with the ground truth unit and plotting at the point (0, 1). All remaining units would plot along a horizontal line near a FD rate of 100%, varying along the horizontal axis only insofar as their average templates happen to have the same shape and spatial distribution as the ground truth waveform.

Units near the center of the scatterplots in Figures 6b-e indicate spike sorter confusion. A relatively lower FD rate means that a unit has a high proportion of its spike events in close temporal proximity to ground truth spike events. Relatively higher template projections on the ground truth template indicate units whose average spike waveform is similar to the ground truth. Together, these two pieces of evidence imply that the unit includes a high proportion of spike events that are the direct result of detecting ground truth spikes. Units near the center of these scatterplots are thus likely to be false units that result from mixing ground truth spikes with either noise or spikes from other neurons.

Analysis of the templates for the units discovered by FBP confirms the impression that it split the ground truth neuron. The two best matching units have low FD rates and high ground truth template projections (Figure 6b, lower right). The templates for these two units bear a strong resemblance to the ground truth (Figure 6g, top 2 examples). The top template has a distinctly smaller amplitude and reduced hyperpolarization phase compared to the bottom template, causing FBP to split them into two separate units. Of the 22 units discovered by FBP, the worst is indicated by the arrow in Figure 6b. It had a relatively small number of spikes, but about 37% of them corresponded to ground truth spikes. This unit’s template (Figure 6g, bottom example) bears some resemblance to the ground truth template with a much smaller amplitude: it is likely a mixture containing some ground truth spikes and some smaller events.

The template analysis agrees that MountainSort4 split the ground truth neuron into 2 units with a relatively large number of spikes and a third unit with a small number of spikes: all 3 units plot in the lower right of the scatterplot (Figure 6c). Their templates (Figure 6h) all resemble the ground truth template and the bottom one indicates that the unit with a low spike count arose from alignment errors. For SpyKING-CIRCUS (Figure 6d), unit 0 plots perfectly in the lower right corner of the graph. One other unit lies toward the center of the scatterplot (Figure 6d, arrow) and has a template that bears some resemblance to the ground truth template (Figure 6i, top example). However, its template has a rather small amplitude (note that scale bar change in Figure 6i, j), suggesting that the unit may be a mixture of noise with occasional ground truth spikes. For KiloSort2 (Figure 6e), unit 0 plots perfectly in the lower right corner of the graph, and several units plot near the center of the scatterplot, including one (Figure 6e, arrow) with nearly the same number of spikes as the ground truth unit, about 50% FDs, and a projection that is about a 46% match for the ground truth template. The other 2 example templates (Figure 6j, bottom examples) suggest similar errors, but with much smaller overall amplitudes.

Finally, we note the differences in the number of units found by the different sorters: FBP, 22 units; MountainSort4, 81 units; SpyKING-CIRCUS, 56 units; KiloSort2, 116 units. The small number of units found by FBP reflects an intentional philosophy. FBP is conservative about identifying units, avoids placing spikes that actually are noise into units, and has an automated procedure to delete likely false units. The other sorters identify many units; a few plot in the middle and most plot in the upper left corner of the scatterplots in Figures 6c-e. Many have an extremely high number of spikes suggesting that they represent large mixtures of multi-unit activity and noise. The output of all sorters on this dataset would require further post-processing to correctly identify well-isolated single neurons.

### Identifying sorter errors in post-processing

#### Strategy

In the absence of ground-truth data, can we assess the quality of a sort, decide whether a neuron was split into pieces, and identify putative units that actually are contaminated by noise or multi-unit activity? Attempting to manually repair the individual units output from a spike sorter for multichannel datasets with multiple neurons is virtually impossible. The sorting algorithms work in a higher dimensional space and objectively consider far more information than any human could for every spike detection and assignment. Instead, we advocate simple decision-making rules for post-processing. After sorting, we eliminate units whose SNR is too low, or number of inter-spike intervals (ISI) less than an absolute refractory period are too high, or have too few spikes to be experimentally considered. Then, we compare the templates, autocorrelograms (ACGs), and CCGs of all remaining units pairwise with one another and make a simple decision. Either (1) the two units show evidence of being the same unit, and should be combined, (2) one of the two units shows evidence of being a mixture of the other and should be deleted, or (3) the two units are legitimate and different. We show 2 examples of how this could work, one for the ground-truth recording analyzed above (Figure 6), and one for a second ground-truth recording.

#### Kampff2015-09-03-Pair-90B (Figures 6, and 7)

The ACGs of FBP units 0 and 1 (Figure 7c, d) have zero refractory period violations (empty bins at t = +/- 1 ms) indicating that both are well isolated. Combining units 0 and 1 yields an ACG without refractory period violations (Figure 7b) as would be expected from two pieces of the same neuron. The CCG between units 0 and 1 (Figure 7e) shows that unit 0 has a large number of spikes at short ISIs of about 3-4 ms after spikes of unit 1, consistent with unit 0 corresponding largely to the second spike of bursts in the ground truth neuron. The same CCG does not show a feature we would expect if there were different neurons, namely overlapping spikes at time bins from −2 to 2 ms. It also does not show the sharp peak we would expect at zero if the two units shared some spikes. Returning to the templates for units 0 and 1 in Figure 6g shows that they differ subtly, as might be expected of the first and subsequent spikes in a burst. Even without ground-truth data, consideration of the templates, the individual ACGs, the combined ACGs, and the CCGs for each of the 22×22 pairs of units leads to the conclusion that one of the neurons in the recording was split between units 0 and 1. Comparison of the combined ACG of units 0 and 1 (Figure 7b) to the ground truth ACG (Figure 7a) confirms that this would have been a good decision.

**Figure 7.**
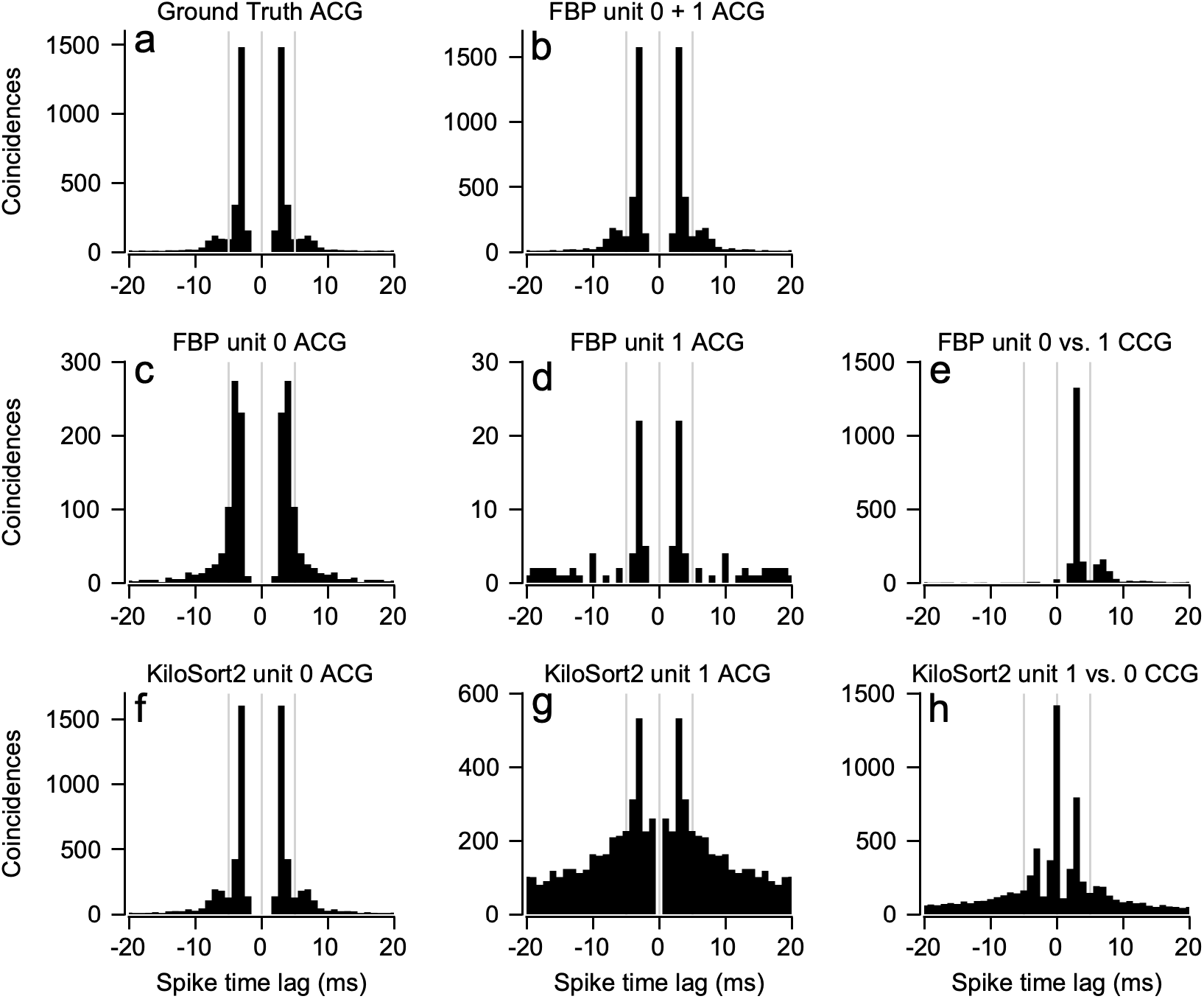
Post-processing analysis of the units discovered by FBP and KiloSort2 in the SpikeForest Kampff 2015-09-03-Pair-9-0A dataset. ACGs and CCGs are derived from the sorts included in Figure 6. ***a)*** ACG of the ground truth spike times. ***b)*** ACG of a unit made by combining the spike times of FBP units 0 and 1. ***c-d)*** ACGs of FBP units 0 and 1. Note the change in scale of the y-axis. ***c)*** CCG of the spike times of FBP unit 0 with respect to the spikes on unit 1. ***f-h)*** Same as *c-e* but for KiloSort2 units 0 and 1. Unit numbers correspond to the units from Figure 6a.

The same post-processing strategy for KiloSort2’s units 0 and 1 suggests a different, but equally good decision. The ACG for unit 0 (Figure 7f) is clean. The ACG for unit 1 (Figure 6g) shows evidence of refractory period violations, with many ISIs shorter than 1.5 ms (bins at t=-1 and t=1). The CCG between unit 1 and unit 0 (Figure 7h) indicates that one of the two units is likely a mixture of the other because there are a large number of spikes with identical event times. Without ground-truth data, we would conclude that unit 1 is probably contaminated with multi-unit activity and/or noise. We would choose to delete unit 1 and keep unit 0. Comparison of the ACG for unit 0 (Figure 7f) with the ground truth ACG (Figure 7a) confirms that this would be a good decision. FBP and KiloSort2 find different paths to the same correct conclusion.

#### Kampff C26 (Figure 8).)

Based on their templates, KiloSort2 and FBP both discover units 0 (Figure 8c, blue) and 1 (Figure 8c, orange) with very different templates that appear to be consistent across sorters. For both sorters, there are a small number of refractory period violations in the individual ACGs of units 0 and 1 (Figure 8f, g, m, n), their combined ACG (Figure 8h, o) shows elevated refractory period violations, and the CCG between them (Figure 8i, p) does not show an implausibly large number of simultaneous spikes at time 0. Based on these measures, all available in the absence of ground-truth data, we would conclude that they are different neurons with a tendency to fire together in a manner typical of neighboring neurons. For FBP, we would accept unit 0 because it is well isolated, without knowing that it has missed 23% of the ground truth spikes (Figure 8a, black bars). For KiloSort2, we would accept unit 0 because it too appears well isolated, in spite of the fact that 12% of its spikes are not from the ground truth neuron (Figure 8a, green bars). We would similarly accept unit 1 for both sorters but the ground truth in this case is not known. Scatterplots of FD rate and projection on ground truth template (Figure 8d, e) confirm that KiloSort2 and FBP did a good job of separating the ground truth unit from other neurons in the recording.

**Figure 8.**
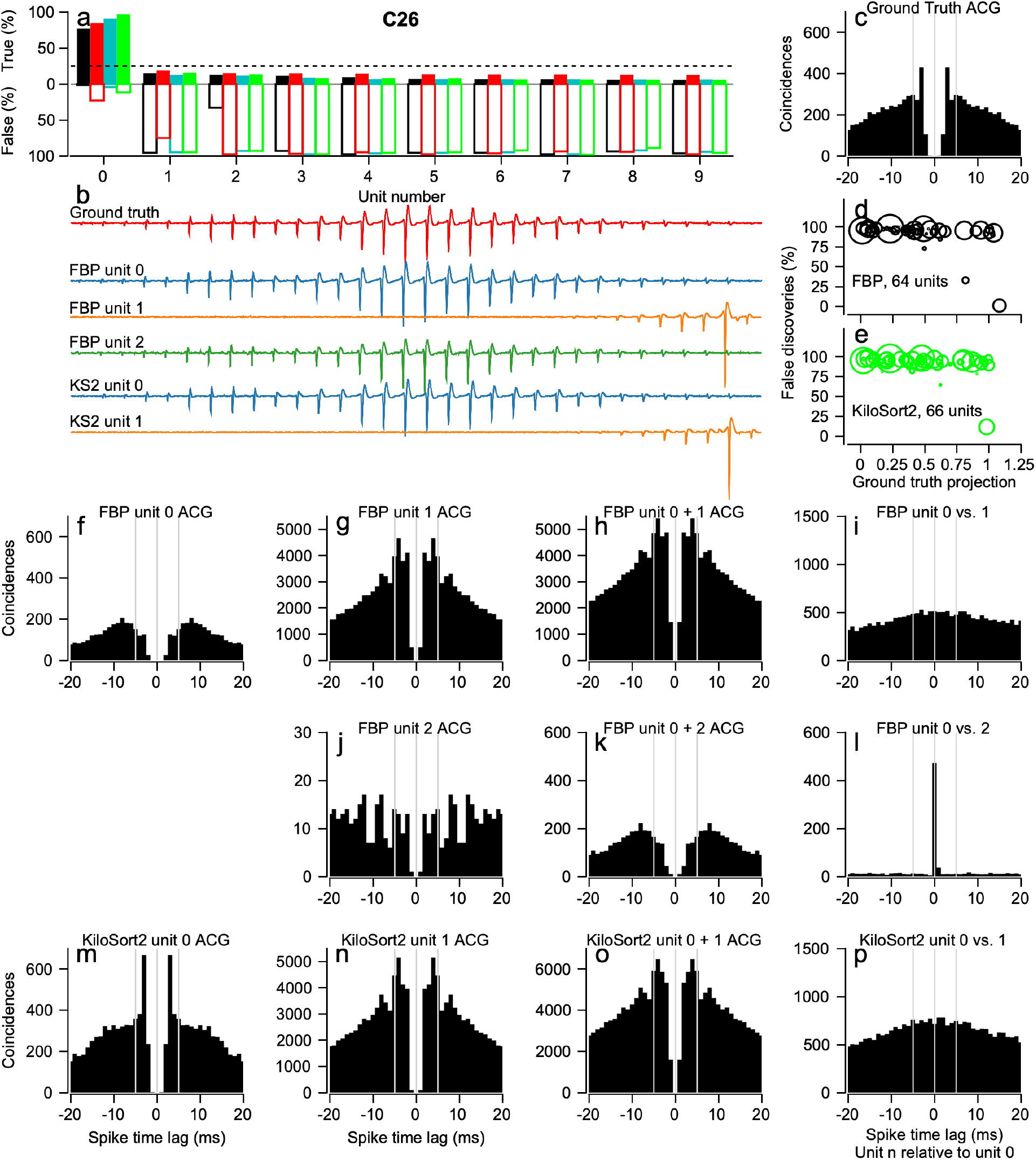
Use of post-processing to evaluate the quality of spike sorter performance for the SpikeForest Kampff c26 ground truth dataset. ***a)*** The graph plots the TP and FD rates ordered by TP rate for the 10 units with highest TP detections of the ground truth unit. Bars above the abscissa (filled) show TP rates, those below the abscissa (unfilled) show FD rates. Data are shown for each of the 4 sorters tested. FBP, black; SpyKING-CIRCUS, cyan; MountainSort4, red; KiloSort2, green. ***b)*** The traces illustrate example templates for units chosen for additional analysis. ***c)*** ACG of ground truth unit. ***d-e)*** Scatterplots of FD rate vs. normalized template projection on the ground truth template for all units output by FBP and KiloSort2. The actual ground truth unit (not shown) would plot at (0, 1). The area within each symbol is linearly related to the number of spikes discovered for its corresponding unit. ***e)*** CCGs and ACGs for units with the top TP rates. Ground truth unit SNR for the ground-truth neuron in this data set is 3.67 on channel 16.

FBP also outputs a unit 2 with a template very similar to that for unit 0. The ACG for unit 2 (Figure 8j) has few refractory period violations, as does the combined ACG for units 0 and 2 (Figure 8k). The CCG between units 0 and 2 (Figure 8l) shows many coincidences, suggesting that unit 2 is largely redundant with unit 0. Given that unit 2 has few spikes that are not already in unit 0 and few spikes overall, we would consider it a strong candidate for deletion. The number of FDs in the ground-truth analysis (Figure 8a, black bars) and this unit’s position near the center of the scatter plot (Figure 8d), supports deletion and suggests that unit 2 is a false unit made up of a mix of ground truth spikes and noise or spikes from other units.

The ground truth unit of file C26 has the lowest SNR of any of the ground truth examples. Consequently, FBP misses spikes, and KiloSort2 adds FDs. SpyKING-CIRCUS discovers an in-between result of 91% TP rate, 4% FD rate, and no additional units with suspiciously high TPs (Figure 8a, cyan bars). The C26 dataset exemplifies how different sorters can have different strengths and weaknesses, and that some datasets might not be sorted ideally by any algorithm.

## Discussion

*Caveat emptor* should be the motto of anyone who embarks on spike sorting. We analyzed the sorting quality of 4 different algorithms on contrived synthetic datasets and actual neural recordings from multi-contact arrays with ground truth. We conclude that no current spike sorter is definitively the best in all situations. All sorters exhibit failure modes and each has its own strengths and weaknesses. Because of the tendency of sorters to discover false units that are either multi-unit activity or a combination of a single neuron and noise, the output of each sorter tested requires automated and/or manual curation if the research goal is to have well isolated single neurons. Our new sorting algorithm, FBP, was designed to avoid specific failure modes that are critical to our scientific objectives, and to enable a rational and feasible post-processing strategy.

The broad rule for choosing a spike sorter should be its performance on a particular application. We require resolution of overlapping spikes from single units to perform CCG analyses on data collected from low-density 16 to 32 contact recording probes with nearby neurons firing spikes at rates as high as 100 spikes/s in baseline conditions. For these purposes FBP is the best choice. It resolves pairs of spike overlaps optimally and is conservative enough to minimize false units arising from multi-unit activity or contamination of a single unit with noise. FBP’s conservatism discovers a small enough number of units to enable manual curation using ACGs and CCGs. Other spike sorter use cases may vary in tolerance for recordings of multi-unit activity versus isolation of single neurons, resolution of overlapping spikes, contamination by noise, and other factors.

We learned that FBP was ideal for our use case through (1) careful design and a full understanding of the sorting algorithm and (2) tests using synthetic data that mimicked our use case. The other 3 sorters we tested have been similarly demonstrated to perform well in their designed use conditions where FBP might show weaknesses. The spike sorting challenge is that the number of variables present in real neural recordings creates an infinite set of possible test cases and failures dependent on a specific experiment, e.g., the number of channels, their proximity to one another, the density of neurons, neuron firing rates, temporal/spatial overlap of neighboring neurons, and similarity between neurons’ templates, to name a few common characteristics to consider. Thus, we advocate that the best sorter be chosen by testing on realistic use cases with synthetic or ground-truth data.

The known advantages for the sorters we tested appear to be features specifically designed into them. Some are designed to sort quickly and produce a large number of units, while FBP was designed to be conservative and deal with overlapping spikes and drift in space and time. We created FBP to perform consistently across single electrode recordings, tetrodes, as well as low-density and high-density arrays. We demanded that given sufficient SNR, and 2 discriminable neurons, FBP could sort the units perfectly in synthetic data for 1 to 8 channels. We included a noise criterion to reduce the number of false units. The penalties for these decisions are slowness of sorting and a high threshold for spike detection. Both KiloSort2 and SpyKING-CIRCUS are aimed at sorting numerous channels from high-density arrays and are capable of sorting these types of datasets much faster than FBP but are less strict in their output units. KiloSort2 was designed to sort data from NeuroPixels and, not surprisingly has occasional problems sorting lower channel count, low density datasets.

Finally, we have shown that a rigorous post-processing step is a necessity if the goal is to obtain single-unit recordings. Refractory period violations in ACGs reveal whether a unit includes multi-unit activity or contamination by noise. A high probability of coincident spikes in CCGs reveals units that share spikes, implying that they should be combined or one should be deleted. Combined ACGs between pairs of units provides evidence about whether they should or should not be combined into one unit. Post-processing is not fool-proof, but we have provided some examples where it can lead to a good decision in the absence of ground-truth data. We designed FBP with these post-processing steps in mind because post-processing scales poorly with increases in the number of units discovered by a sorter. With 20 units, post-processing could involve as many as 400 CCGs. With 100 units, inspection of 10,000 CCGs is implausible. To us, though, the effort to perform rigorous post-processing seems valuable to ensure the validity of neuroscience data derived after spike sorting.

## Methods

### Overview

Our data comprise an nC-channels by nS-samples matrix of the continuously sampled and digitized raw recorded voltage. Our analysis has two phases that operate more or less independently. <u>First</u>, we perform multi-step dimension reduction to create an average template that concatenates waveforms across channels to identify uniquely each neuron in the recording. The steps are: (1) filter the voltages and divide a long recording into shorter segments so that the voltages are quite stable in space and time; (2) analyze each channel separately for each segment to identify clips of voltage that cross a threshold and gather the clips into clusters of like waveforms; (3) condense across channels, remove duplicates, and average across clusters of clips to define a set of template waveforms, concatenated across channels, that define all the neurons in the segment. <u>Second</u>, we abandon the clips and work from the concatenated-template waveforms, stepping through the voltage traces across channels to identify waveforms that statistically are assigned to the neuron represented by each concatenated-template. The steps are: (1) perform “binary-pursuit” template matching on the segment to identify voltages that belong with each neuron’s concatenated-template; (2) disambiguate overlapping waveforms and define the spike times of each waveform; (3) stitch the data back together across all segments to define the total set of recorded neurons. We explain the details in separate sections below.

### Data preprocessing

We apply a zero-phase bandpass digital filter to each channel in the raw voltage matrix. Then, we break the voltage into segments of 5-minutes duration, with each segment overlapping the previous segment by 2.5 min. Sorting spikes in segments provides computational speedup: our approach depends on initial over-clustering, over-clustering in longer segments creates more clusters, and our algorithm runtime increases with the square of the number of clusters. The use of reasonable-length segments with temporal overlap between adjacent segments also enables strategies to compensate for drift in the waveform shape or amplitude of individual spikes across time and for spatial drift across channels on a multi-contact probe.

Next, we find a threshold (*Th_c_*) for event detection on each channel *c* as:

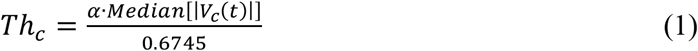

Here, the “median absolute deviation” of the time-varying voltage (*V_c_*) on channel *c* over time is divided by 0.6745 to provide an estimate of the standard deviation of the voltage that is robust to outliers. The factor α scales the number of standard deviations to use as the threshold. We then define the SNR on a channel as the maximum minus the minimum voltage on that channel divided by *Th_c_*.

We remove the correlations between channels by spatially whitening the data using a “zero phase component analysis sphering” (ZCA) transform. We estimate the covariance matrix for the ZCA transform for each pair of channels by randomly sampling 10^6^ data points from the ‘noise’ by choosing voltage points that are within +/- α*Th_c_* on both channels. We compute the inverse correlation matrix and multiply it by the entire segment voltage matrix to obtain the ZCA transformed voltage. The ZCA-transformed voltage is thus whitened and has a channel-wise covariance matrix for subthreshold datapoints that is the identity matrix. All subsequent analyses are performed on the ZCA-transformed voltage, which we will henceforth simply refer to as the voltage.

### Event detection

Our first spike-sorting step breaks the voltage into “spike-event clips” that will, in the end, define the characteristic shape of the voltage deviation associated with each particular neuron’s firing event. Each clip has a specified duration encompassing a window of time centered on a spike event. The goal of an ideal clip width is to capture the full characteristic deviation of a spike event without being long enough to exceed the unit’s absolute refractory period. To define all clips, we recompute *Th_c_* for each channel from the whitened data using Equation (1). We find all positive or negative deflections of the voltage trace that cross plus-or-minus a*Th_c_*, and we identify the local absolute maximum or minimum within one clip width of each threshold crossing. This local extremum is selected as the tentative spike event time, and we extract clips centered at this time point. We ignore any threshold crossings that follow a spike-event clip by less than a clip width, so that each spike-event clip indicates a local positive-negative (or negative-positive) excursion that is likely to correspond to a single action potential. We assemble the full set of spike-event clips by extracting snippets of the voltage traces from *cw_pre_* samples before to *cw_post_* samples after each putative spike event time, where *cw_pre_* + *cw_post_* + 1 = *cw_total_*, the total number of time samples comprising each clip.

We refine the temporal alignment for each spike-event clip using a Mexican hat wavelet because our initial, tentative, choice of the time of a spike relies solely on local extrema, which are susceptible to outliers due to noise. Moreover, because of the choice of the absolute extreme voltage as the time of a clip, spikes that are roughly equal in positive and negative going magnitude will be aligned arbitrarily on their peak or trough in different instances due to random fluctuations. To correct this, we compute the cross correlation between the clips and Mexican hat wavelets of a series of frequencies and find the frequency that gives the maximal absolute (positive or negative) cross correlation for a given clip. We then adjust the spike-event time for the clip so that it is aligned optimally with the wavelet and resample the spike event clips to be centered on the newly adjusted spike-event times. The cross correlation gives a better measure of spike timing that takes all data points in the voltage clip into account, rather than noisy local extrema. We find that extra care in choice of spike-event time is critical for accurate principle components analysis and clustering in subsequent steps.

Now that the clips are optimized on the channel we are sorting, we collect the simultaneous voltage clips from across all nearby channels in the current channel’s neighborhood, defined as the set of channels that are user-specified to account for the brain region of recording and the electrode configuration of the recording probe. We concatenate the clips from all individual neighborhood channels to form a single multichannel clip for each spike event (Figure MF1, top center). We then remove all clips that have their absolute maximum value on a different channel than the one we currently are sorting. We do so on the assumption that they belong to a different set of neurons and will be sorted when their respective channel is being considered. Later, we address the exception to this assumption, where an individual neuron has a variable spike amplitude that causes it to appear with a maximum that varies across neighboring channels in different clips.

### Clustering

We identify the neurons present in a recording and their individual waveform shapes by an elaborate, multi-step procedure. We initially cluster the clips defined only on the channel we are currently sorting. At the end of clustering, we use the concatenated multi-channel clips to take advantage of any additional information afforded by the spike waveform shapes across all channels in the neighborhood. To specify the space to be used for clustering, we use principal component analysis (PCA). We select principle components (PCs) based on their reconstruction accuracy, where the goal is to identify the minimal number of PCs that maximizes the average clip reconstruction (Abdi and Williams 2010). Briefly, we compute the PCs across our matrix of spike-event clips, *X*. For each of the identified PCs, we compute the mean squared error between the clips and the reconstruction using solely the current PC. The reconstructed spike-event matrix, *M*, is computed as a linear model of *X* and the PC matrix, *Q* as:

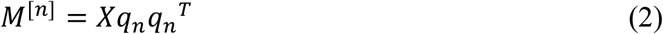

where *M^[n]^* is the reconstruction derived from the *n_th_* PC and *q_n_* is the *n_th_* PC column vector of *Q*. The reconstruction error for the n^th^ PC, *RE_n_*, is computed as the mean sum of squared error between the original spike-event clips and their PC reconstruction:

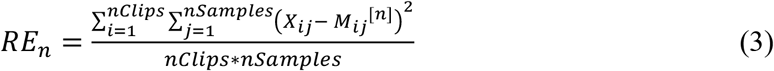

where *nClips* is the total number of spike event clips (rows in *X*) and *nSamples* is the total number of samples in each clip (columns of *X*). We then reorder the PCs from smallest to greatest reconstruction error. Note that this method deviates from the standard method of ordering PCs by the eigenvalue “variance accounted for” (VAF). Ordering the PCs based on eigenvalue-based VAF serves to maximize the variance across clips rather than the variance within clips. In instances where there is substantial noise across clips, eigenvalue-based VAF ordering of the PCs may result in a choice of PCs that overfits the noise. In contrast, ordering the PCs by reconstruction accuracy more strongly weights the within clip variance, which generally corresponds to the waveform characteristics of individual neurons’ spikes.

**Figure MF1.**
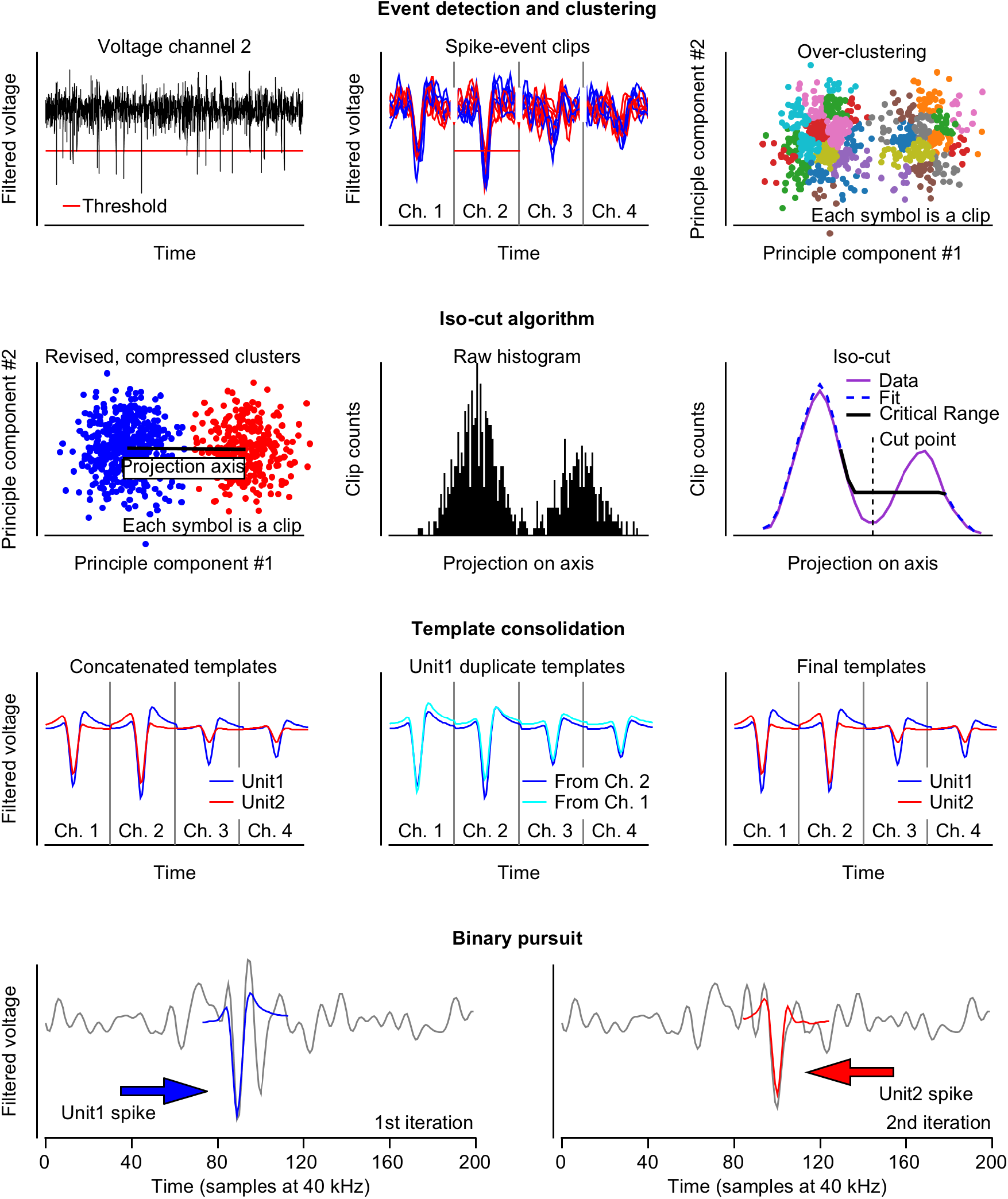
Details of sorting procedure for each step of spike sorting pipeline for artificial tetrode dataset. *First row: detection of spike events, clips and clustering space*. (*left*) Filtered and ZCA transformed voltage example for channel 2. Horizontal red line shows the spike event detection threshold. (*middle*) Spike clips for threshold crossing events. Clips are taken separately for each of the 4 channels and concatenated across all channels. Only a subset of clips is shown for clarity. (*right*) Principle component (PC) projection of all spike clips onto the first 2 PCs. The clustering algorithm is seeded by overclustering (colored groups). ***Second row: isocut algorithm*.** (*left*) Clusters are compared pairwise with their nearest neighbor (final comparison shown). Data for each cluster are whitened and projected onto the line connecting their average centroids (black line). (*middle*) The projected PCs for the pair are converted into a 1D histogram. (*right*) The actual data histogram is smoothed (magenta) and a null distribution is formed using unimodal isotonic regression (orange). The data and null distribution histogram bin counts are compared within the critical range (black) using a multinomial goodness of fit test. ***Third row: template consolidation across channels*.** (*left*) Ground truth templates used to create the artificial dataset. Templates represent the average spike waveform for each unit. Note that both have maximal value on channel 2. (*middle*) Due to noise, clips for unit 1 sometimes have maximal deviation on channel 1, and other times on channel 2. Because data are sorted independently for each channel, the clustering algorithm outputs 2 different templates, both of which correspond to unit 1. (*right*) The spike clips corresponding to unit 1 on channel 1 and channel 2 merge and so they are combined to form a single template for unit 1. ***Fourth row: binary pursuit template matching. Only data for channel 2 are shown for clarity*.** (*left*) First iteration of binary pursuit. Voltage trace (gray curve) showing overlapping spikes from unit 1 (blue arrow) and unit 2 (red arrow). Binary pursuit finds a maximal match for unit 1 template (blue curve) and adds a spike event to unit 1. (*right*) Second iteration of binary pursuit. The voltage trace (gray curve) is the residual voltage trace from iteration 1. The template of unit 1 was subtracted during the first iteration, leaving behind only the spike waveform of unit 2 (red arrow). Binary pursuit correctly identifies and assigns the spike from unit 2 (red curve).

After ordering the PCs, we determine the optimal number of PCs that should be used to subsequently cluster the spike waveforms. We iteratively add the reordered PCs one at a time to the linear reconstruction model and compute the residual error at each step.

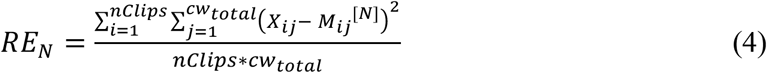

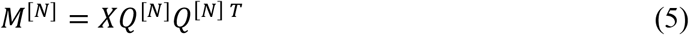

Here *M^[N]^* is the reconstruction derived from the first *N* PCs, and *Q^[N]^* represents the first *N* columns of the matrix Q. This yields the corresponding residual error *RE_N_*, which is the residual of the reconstruction based on the first *N* PCs. From *RE_N_* we define a function quantifying the change in the mean residual error after the addition of each PC, computed from the ratio of the residual error between steps as:

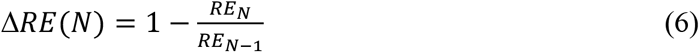

Δ*RE*(*N*) quantifies the relative improvement in the residual error of the reconstruction by the addition of the *N^th^* PC to the model. We define *RE_0_* by computing the residual error with respect to the mean value over all *X_ij_* in *X*. Local maxima of Δ*RE* indicate points where addition of a PC to the reconstruction provides no improvement. We choose the basis of our clustering space to be the first *k* PCs, where Δ*RE*(*k*) is the first local maximum of Δ*RE*.

Next, we project all spike-event clips from the channel being sorted onto the *k* PCs chosen as the clustering space above and begin a clustering procedure similar to that proposed by Chung et al. 2017. We begin by over-clustering the data (Figure MF1, top row). We start with the full set of clips in PC space and follow an iterative strategy to select a point as a new cluster center, and then add a cluster by reassigning all spike-event clips to their nearest cluster centroid. Our strategy uses the k-means++ algorithm, which randomly selects a point in the data with a probability that is proportional to the distance of each point to its current centroid (Arthur and Vassilvitskii 2007). We complement this approach with a deterministic cluster seeding that always chooses the data point at the 95^th^ percentile of distances from its cluster center. We continue the over-clustering process until the median cluster size is the smaller of 100 or the number of clips divided by 1000. At this point, we have many more clusters than unique waveforms in the original recording.

The next step is to determine how the clusters should be separated and combined on each individual channel (we combine across channels later). We mimic the central concepts of the isocut algorithm (Chung et al. 2017) but with key differences in smoothing and statistical tests. The main idea is to assess the degree of similarity and difference pairwise by considering each pair of clusters that are mutually closest to one another, measured as centroid to centroid distance, and asking whether the two clusters should be merged or split. We first whiten the pair and project their points onto the axis that connects the two centroids to obtain a decorrelated, 1-dimensional representation of the clusters to be able to capitalize on low dimensional statistical analyses (Figure MF1, second row). Given the 1-D distribution of data points for each pair of nearest neighboring clusters, we ask whether the distribution is consistent with our null hypothesis of a unimodal distribution, or whether it is multimodal and should be split into separate clusters. We smooth the distribution using a non-parametric kernel density estimator that is robust to multimodal distributions (Botev et al. 2010) and therefore substantially improves the ensuing fits and statistical tests. We treat the smoothed distribution as our ‘observed’ data and derive all subsequent analyses from it.

To decide whether to separate or combine two distributions, we obtain a null hypothesis for their combined distribution by fitting the observed data with an optimal unimodal distribution via isotonic regression (Figure MF1, second row right panel, dashed blue curve). We choose a critical region of comparison between the unimodal and observed distributions as a subset of the two distributions such that (1) the difference between the null and observed distributions is maximized and (2) the first and last points are identical for both distributions, or lie at the peak of the unimodal distribution (Figure MF1, second row right panel, black). Intuitively, this means finding a window that lies to the left or right of the unimodal peak where the observed data dips below the unimodal fit maximally for consecutive observations. Such regions are easily discovered, if they exist, due to the monotonicity of the isotonic fit.

We use a multinomial goodness of fit test to compare the null and observed distribution bin counts within the critical range, instead of the score suggested by Chung et al. (2017), because we found it to be adequately but not overly sensitive. The multinomial goodness of fit test can be computed exactly for relatively small sample sizes. For larger samples, we performed a permutation test version. If the statistical test yields a p-value greater than a user defined threshold (e.g. p = 0.05), then we accept the null hypothesis and combine the two clusters. If the p-value is less than the criterion, then we separate the two clusters at a value chosen as the peak of the isotonic regression fit to the difference between the null and observed distributions within the critical range (Figure MF1, second row right panel, vertical dashed line). The peak of this fit indicates the maximal deviation of the observed distribution below the null distribution. Data from the two clusters are reassigned on each side of the cut point and thrown back into the clustering algorithm. We repeat the entire pairwise comparison of mutually closest cluster pairs until no additional merges occur. The remaining clusters form a first hypothesis for the set of individual neurons on the sorted channel.

Next, we refine our clusters further. We compute a template, defined as the average clip waveform for each cluster, align each individual clip in the cluster on the average template by centering the index of their maximal cross correlation, and recompute all clips by shifting their time to be aligned on their newly aligned event times. Within each cluster, we perform “branch-PCA” (Chung et al. 2017) by repeating the clustering procedure described above. Because the PCs are now are calculated within each cluster instead of across all clips, we obtain a different PCA space that may be more sensitive to differences between clips within each cluster: this step can only increase the number of clusters. Finally, we perform the branch-PCA procedure on the multichannel clips created by concatenating the voltage clips across all channels in the current channel’s neighborhood. Differences in waveform voltage traces across channels further separate out individual neurons, resulting in our final set of neurons on the current channel.

### Extracting and consolidating templates

Next, we consolidate the clusters, which were independently sorted one channel at a time, into a refined set of putative neurons across all channels, still working one 5-minute segment of data at a time. Consolidation involves two considerations.

The first consideration is whether a cluster may represent spikes from 2 separate units that overlap with each other reliably enough to form their own cluster. Assume that clusters A and B each have characteristic waveform shapes, and a third cluster C has a waveform that is a sum of the waveforms A and B. To assess this possibility, we iteratively assign each cluster in the channel to be “C” and compute a “template” defined as the average clip for C across all channels in the recording, regardless of the input neighborhood. We then ask whether we can shift and add a combination of any two other templates, A and B, to account for > 75% of the sum of squared variance of template C. If so, we no longer consider cluster C to be a candidate to be a separate neuron because it is either (1) a combination of A and B or (2) so similar to a combination of A and B that our algorithm will not reliably discriminate it. We therefore remove cluster C and associated data from any further consideration. To eliminate C as a separate neuron, we additionally require that:

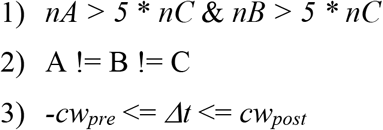

where *nX* indicates the number of clips in cluster X, and Δ*t* is the shift between A and B for their sum to be compared to C (derived from the clip width). Note that removing cluster C does not eliminate its clips from consideration as spikes from neurons. They may be assigned to neurons in the later, template matching stage of sorting.

The second consideration is whether a single neuron has given rise to clusters on more than one channel. Due to noise, spike variance, amplitude drift and other intangible factors, the spikes from a single neuron could sometimes be maximal on channel N, and other times on channel M. Because we sorted N and M independently, we could have two copies of the neuron, one on each channel. Working with templates that are concatenated across all channels in the recording, we search iteratively through the average templates pairwise for all pairs of clusters from individual channels. For each pair, we find an optimal time-shift (Δt) based on the cross correlation in the concatenated templates and compute the Euclidean distance, *d*, between average templates (*T_i_*) at that time shift:

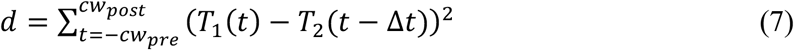

For unrelated pairs, *d* will be large; for pairs that share clips from the same neuron, *d* will be small because their templates are very similar. Optimal Δt > 0.5 * *cw_total_* are ignored. For the pair with the smallest *d*, we then create a more meaningful set of channels to consider based on the signal-to-noise ratio (SNR) in the templates (*T_c_*) for each individual channel. We define SNR for each channel as:

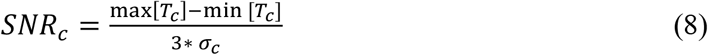

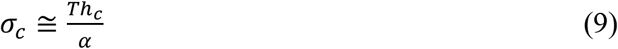

where *σ_c_* is the estimated standard deviation of the noise and *Th_c_* and *α* are from Equation 1. We use the optimally-aligned clips to create a new set of concatenated clips only for channels where *SNR_c_* for at least one of the units is greater than 0.5. We now have two new clusters, and we test whether the two clusters merge using the clustering algorithm described in the previous section. Here, we use a 2-dimensional space by projecting all clips onto the average templates for the two clusters being compared rather than a PC based space. If the clusters merge across channels, they are considered to come from the same neuron and their spike-event clusters are combined. If they do not merge, the clusters are considered to be distinct and are not checked further against any other clusters. We repeat the merging procedure iteratively until none of the remaining clusters merge with their nearest template pair. This leaves us with the spike-event clips for a set of unique neurons that respects interactions across all channels.

### Assigning spike times and unit labels: modified binary pursuit

At this point, clustering has identified a set of clips corresponding to each putative neuron in the recording. To identify the spike event times for each neuron, we compute the average template concatenated across all channels for each neuron and use template matching to assign all spikes in the voltage trace to the appropriate unit. In effect, we now discard all previously found spike event times and clips and start over, but with rich knowledge about the voltage signature across channels of the different neurons in the recording.

We adapted the binary pursuit algorithm of Pillow et al. (2013) to perform template matching. Briefly, their procedure models the recorded voltage traces as a linear combination of the spike waveform templates and noise. They use Bayesian inference to compute the posterior likelihood of the distribution of spikes across space and time, given the recorded voltage and a prior estimate of the probability of detecting a spike from a given neuron (i.e., the firing rate of each identified unit). After some derivation, this means maximizing the log likelihood objective function across all channels in the recording, *L*(*t, T*):

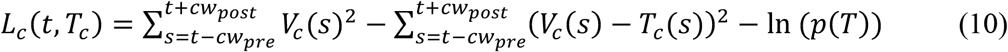

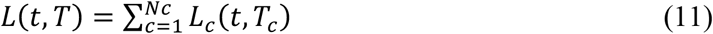

Here *V*(*s*)^2^ is the squared deviation of the recorded voltage at time *s, t* is time with respect to the start of the segment being sorted and *s* spans the clip width centered on *t*. The template, *T_c_*(*s*) is the template of a given neuron on channel *c*, and *p*(*T*) is the Bernoulli probability of observing a spike from the neuron with template *T* at a given time point. Intuitively, the first term of the equation indicates the current sum of squares residual voltage in the window centered at time t, and the second term indicates what the sum of squares residual voltage would become if we were to subtract the template from the voltage.

We do not have an accurate estimate of the prior probability of observing a spike. We therefore chose to drop the prior probability term and instead adopt a noise penalty for each template (*N*(*T_c_*)) that is proportional to the template amplitude of a unit on each channel and the noise on that channel (explained in further detail below). This yields our objective function:

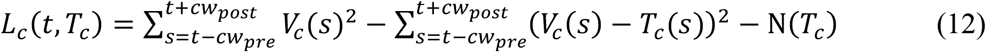

Expanding the *(V_c_(s)-T_c_(s))^2^* term and simplifying yields:

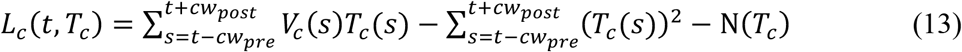

And finally summing across all channels:

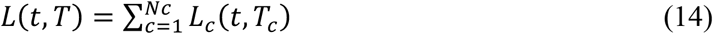

We optimize this function in 3 iterative steps using the greedy binary-pursuit algorithm (Pillow et al. 2013):

1. Compute *L*(*t, T*) for all *t* in *V* and all templates/neurons *T*.
2. If all values *L*(*t, T*) < = 0, then the algorithm has converged, all spikes are assigned to neurons, and we are *done*.
3. For the values of *t* and *T* that maximize *L*(*t, T*)

a. Add a spike event at time *t* to the neuron with template *T*.
b. Subtract template *T* from *V* in the window centered on *t*.
c. Repeat starting from step 1 with the new residual voltage, *V*.

In words, the binary-pursuit algorithm adds a spike at the time point *t* for the neuron with template *T* that maximally increases the posterior likelihood over all *t* and *T*. In our formulation of binary-pursuit without a prior probability, this strategy optimally reduces the residual voltage by subtraction of templates, given that the voltage clip being assigned as a spike exceeds a statistical noise threshold *N*(*T*). Binary pursuit identifies spikes without relying on a single threshold crossing, but rather considering all available data in the voltage trace.

#### Noise bias

The binary-pursuit algorithm requires only that the voltage at any given time is sufficiently similar to a unit template that subtracting it reduces the residual voltage. The algorithm requires noise bias to prevent false positives. The noise bias term is especially important for templates that are low amplitude, or otherwise similar to the noise. Such templates would be repeatedly fit to noise, slowing the algorithm needlessly and contaminating the final set of sorted neurons. The noise term reduces the chance of false positives for well-isolated units, but increases the risk of false negatives for units whose true spiking activity deviates within the range of the noise.

We define the noise bias for each concatenated-template as:

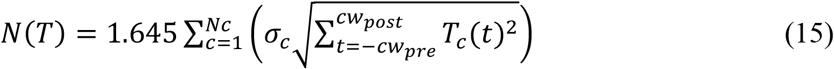

where *N_c_* is the number of channels in the recording, 1.645 is the one-sided 95% critical value for the normal distribution, and σ_c_ is the standard deviation of the noise on channel *c* (Equation 9). Intuitively, the noise term attempts to account for chance variation in our objective function *L*(*t, T*) that is due simply to noise. It estimates the variance of the *V(s)T(s)* term of equation (14) and requires that the criterion to add a spike exceeds the 95% confidence interval of its noise fluctuations.

**Figure MF2.**
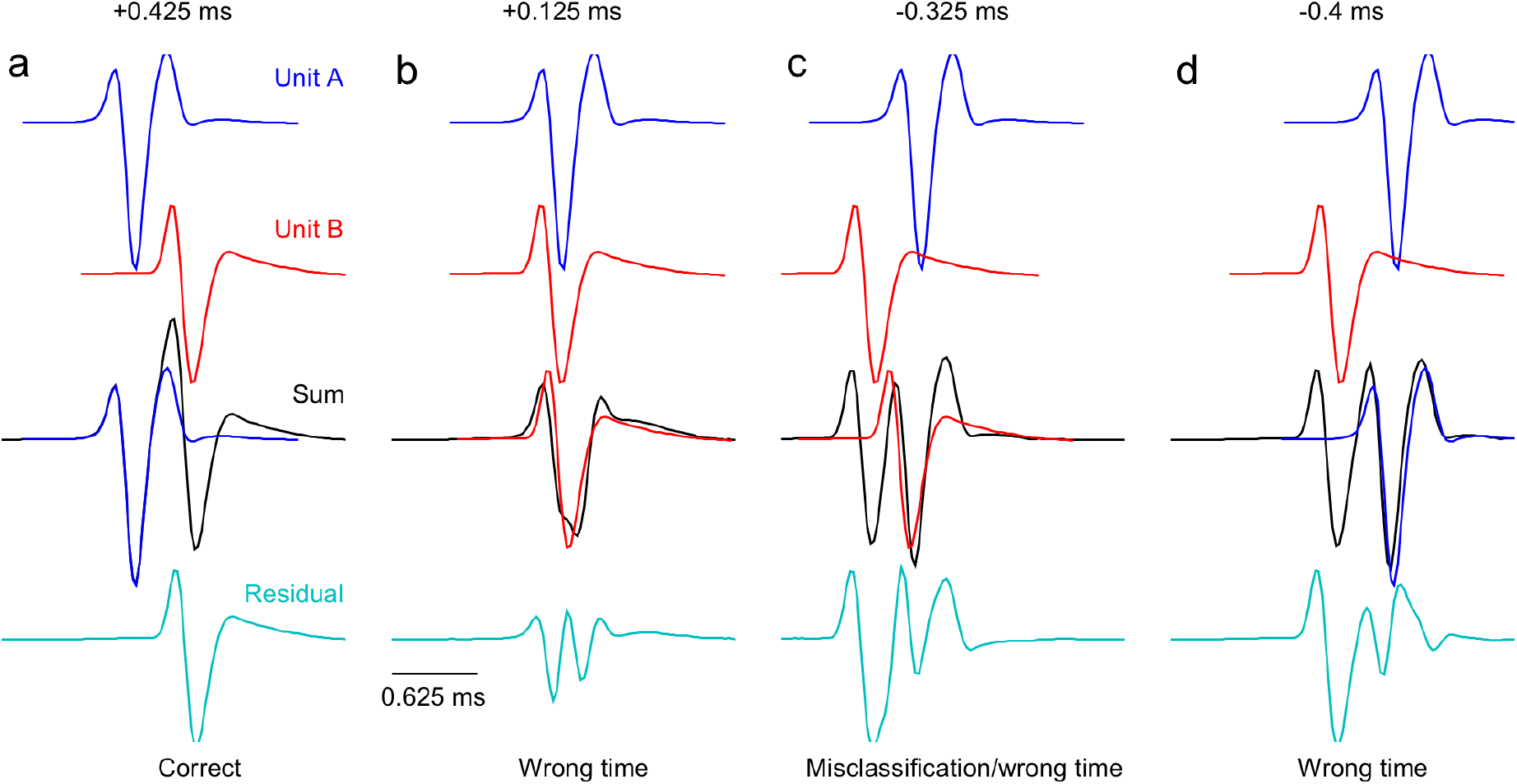
Greedy template matching systematically fails at certain temporal offsets when spikes from two neurons overlap. **Top two rows:** Average spike waveform templates for two units, A (blue) and B (red) at slightly different time lags. **Third row:** A noiseless pseudo voltage trace (black) created by adding the two templates at the temporal offset shown. The binary pursuit algorithm chooses to assign a spike to the unit indicated by the overlaid template. **Bottom row:** The resultant residual voltage after subtraction of the selected template from the original pseudo-voltage. This is the voltage available to the subsequent iteration of binary pursuit.

#### Overlap detection and correction

Optimization algorithms for template fitting, such as binary pursuit, tend to be greedy (Pillow et al. 2013). When spike overlaps are not an issue, greed can help guarantee convergence, but in the presence of overlaps it can lead to deterministic failures.

To demonstrate, we created two single channel voltage templates without any noise (Figure MF2). We added the unit A template (blue) and the unit B template (red) at different time shifts relative to unit A to obtain a pseudo spike voltage trace containing overlapping spikes (black).

- When the two spikes are sufficiently separated in time, binary pursuit is able to accurately identify the two spikes (Figure MF2a). The first iteration of binary pursuit correctly identifies the unit A spike at exactly the correct time (third row) and subtraction of the template from the net voltage yields a residual voltage trace that looks almost exactly like the unit B template (Figure MF2a, bottom row, cyan). The subsequent iteration of binary pursuit easily identifies a unit B spike event, resulting in a perfect sort.
- When the unit B spike follows unit A by only 0.125 ms (Figure MF2b), the summed voltage contains a single, extra wide waveform (third row, black). The first iteration of binary pursuit now assigns unit B at a time point that is incorrect by only 2 samples (red trace overlaying black trace) because subtracting the unit B waveform at that time maximizes the decrease in the residual deviation of the voltage trace. The residual voltage (Figure MF2b, bottom) does not resemble a spike from either unit and does not attract the addition of a spike event during the next iteration of binary pursuit. The unit B spike is detected at a time that was off by 2 samples, but the unit A spike is missed.
- When unit B precedes unit A by 0.325 ms (Figure MF2c), the algorithm assigns unit B at nearly the unit A spike time. The end result is a misclassified spike assignment that is not at the correct spike time for either unit. The residual voltage trace is a mess, and binary pursuit assigns the unit A template at the unit B spike time.
- A larger negative deviation where unit B spikes 0.4 ms before unit A (Figure MF2d) pushes the unit A optimal template match away from the true unit A spike time. The resulting residual voltage, although inaccurate, still allows the second iteration of binary pursuit to correctly identify the unit B spike in the noiseless example of Figure MF2. However, the waveforms clipped out of the voltage at the times assigned to each unit will be poor representations of the true spikes and a realistic level of noise in the recording will increase the risk of total failure.

We amended the binary pursuit algorithm of Pillow et al. (2013) to search specifically for spike waveform overlaps and accommodate the shortcomings of binary pursuit. Our strategy asks, statistically, whether each spike found in binary pursuit belonging to template *T* might have a combination of *T* and another template *A*, such that:

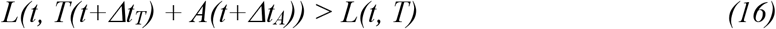

over a range of values of *Δt_T_* and *Δt_A_*; the maximum shift width is limited to one clip-width minus 1. If Equation (16) is satisfied, then we remove the spike that had been created at time *t*, add the previously-deleted template back to the residual voltage, and create a new spike event according to:

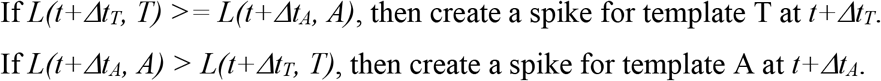

The creation of the new spike causes subtraction of the relevant template from the residual voltage, leaving binary pursuit to find the other spike in the overlap pair in a subsequent iteration.

The benefit of considering whether each spike event is more likely to have arisen from a sum of two overlapping spikes is a guarantee to resolve detected spike events arising from up to 2 overlapping spikes, provided SNR is sufficient and templates are accurate. Our approach can still considerably enhance overlapping spike detection in instances of 3 or more overlapping spikes, but performance may still suffer from the types of deterministic errors shown in Figure MF2. In our own data, instances of 2 overlapping spikes with sufficiently similar waveforms appearing on roughly the same channels is common enough to merit this additional check, while instances of 3 or more overlapping spikes are vanishingly rare.

### Stitching segments

The final step in our algorithm is to clean up the sort for each segment and then stitch together the neurons across segments to obtain a final output that represents each neuron’s spikes for the duration of the recording session. This step takes advantage of the overlap between adjacent segments, where we sorted identical voltages in each channel.

We begin by creating new templates in each segment based on the spike event times assigned by binary-pursuit to the different neurons. We then take three very conservative steps to remove units that are unlikely to represent well-isolated neurons. First, we delete any units whose templates have a SNR less than 1.5. Second, we delete any units with a high degree of multi-unit activity, as ascertained by analyzing their inter-spike interval (ISI) distributions to assess the number of violations of a specified absolute refractory period. We create a histogram of all ISIs less than 500 ms for each unit and compute a “multi-unit activity ratio” as the ratio between the number of ISI counts within the absolute refractory period and the maximum ISI count across all bins. Units are deleted if their multi-unit activity ratio exceeds 0.2. Third, we remove units that appear redundant with other units based on CCGs computed in 1 ms bins for all pairs of units in the sort for the given segment. Iteratively, we find the pair with the greatest number of identical spike event times (bin count at t=0 in the CCG) and ask whether there is sufficient evidence of redundancy that one of them should be deleted. If the jump in the spike count between neighboring bins at t=0 is twice as big as the jump between any other two neighboring bins, then we conclude that the CCG is physiologically implausible and the two units are redundant. We preferentially keep the unit that has the lower multi-unit activity ratio and the template with the larger SNR. If neither unit satisfies both criteria, then we delete them both.

Working with the cleaned-up data, we assign each neuron within a segment to a specific channel, chosen as the channel where that neuron’s template has the highest SNR. Using a concatenated-template based on only the channels within the user-defined neighborhood of the assigned channel, we attempt to match each neuron to itself from segment to segment. We do so in two steps.

First, we test and attempt to take advantage of the assumption that each unit is likely to be stationary on its assigned channel across segments, *S*. We take our complete set of neurons to be the set of neurons present in the first segment. Starting with *S=1*, we proceed in temporal order of the segments, from first to last, considering one channel at a time as follows:

1. Compare all templates in segment *S* pairwise with those for the same set of neighboring channels in segment *S+1*. Using cross correlation between the templates, find the optimal shift that minimizes the vector distance between the pair:

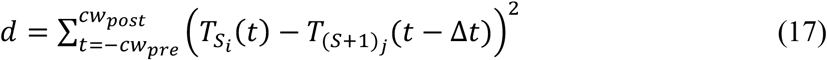 Here *T_Si_* indicates template *i* in segment *S* and *T_(S+1)j_* indicates template *j* in segment *S+1*.
2. For the pair of templates with the smallest distance between them, align the waveform voltages assigned to those neurons according to the optimal shift, truncating as needed.
3. Compute the principal components of the two concatenated-templates (not their clips) and choose the best components defined by the reconstruction error approach of Equations (3)–(6).
4. Project the voltage waveforms from both neurons into this principal component space and define the two neurons as two separate clusters, skipping the initial over-clustering procedure.
5. Test whether the two clusters merge, using the fitting and statistical approach described earlier.

If the two clusters merge statistically, then both are assigned to the neuron in segment *S* and the template in segment *S+1* is removed from any further comparisons.
If the two clusters do not merge, then the template in segment *S+1* is different from the most similar template in segment *S*. It is assigned tentatively as a novel neuron and added to our full set of neurons; the templates from segments *S* and *S+1* are removed from any further comparisons.
6. Return to step 1 and iterate until no more merges occur between any templates in the two segments under consideration.
7. Any further templates in segment *S+1* that did not find a match from segment *S* are assigned as novel neurons and added to the full set of neurons.
8. Increment the segments under consideration and repeat the entire procedure starting from step 1.

Second, we consider whether a unit may have drifted spatially across segments to a new, presumably neighboring, set of channels. We again implement a nearest pairs iterative procedure that advances segment by segment as follows:

1. From the results of the channel stitching above, define “orphan” neurons as all neurons in segment *S* that did not find a match in segment *S+1* plus all neurons in segment *S+1* that did not find a match in segment *S*. The orphan neurons might have drifted across channels on the recording probe between segment *S* and *S+1*.
2. Compute the proportion of spike coincidences, within 1 clip width, between each orphan neuron in one segment and all other neurons in the adjacent segment during the segment overlap time period, when the sorting is based on identical voltages placed in different contexts.
3. Find the pair with the maximal proportion of spike coincidences. If this proportion exceeds 75%, then combine the neurons and removed both from further consideration.
4. Repeat (2) and (3) until no pairs remain with proportion of coincidences above 75%.
5. Proceed to the next adjacent segments and repeat.

As a final step, we remove the duplicate spikes that necessarily are present due to the overlap time between segments. We do so by removing any spikes within a spike width of each other, preferentially retaining the spike that best matches the template. The spike width is computed as twice the interval between the maximum and minimum values of the neuron’s template. We designed the interval that defines overlaps to be small enough so that autocorrelograms remain valid and can be used to identify interspike intervals that violate the neuron’s absolute refractory period, but large enough to account for the fact that waveform shapes and alignments might not be constant across segments and channels.

### Datasets

#### Synthetic datasets

To create synthetic data, we first generated voltage traces of bandlimited noise from 300-8000 Hz. Noise was Gaussian white noise with random phase and scaled to a standard deviation of 0.33, such that 3 standard deviations of the noise was roughly equal to 1 voltage unit. Noise on each channel (*V_c_*) was 30% correlated by creating a shared voltage across channels and an independent noise voltage for each channel separately:

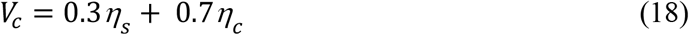

where *η_s_* represents the shared noise across all channels and *η_c_* the independent noise for channel *c*. Simulated spiking events were created by adding templates to the noise. The spike templates were taken from actual recordings, and their waveform shape was identical but multiplicatively scaled separately for each channel as shown in the figures. The timing location of the spike events was determined by a random Poisson process with a refractory period of 1.5 ms or by the spike times output by the leaky integrate-and-fire neural network simulation described below.

#### Neural network simulation

Our simulated feedforward leaky integrate-and-fire network was created using the freely available Brian2 software package (Stimberg et al. 2019). Our feedforward network consists of two units, where the inhibitory unit provides direct inhibition to the second unit, its target unit. The input drive *I*, to each unit was a 5 Hz square wave modulation from 60 to 80 mV that provided constant excitation that kept both units firing at high rates. The voltages of the inhibitory unit *V_i_* and its target neuron *V_T_* were defined by.

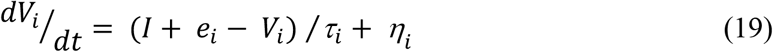

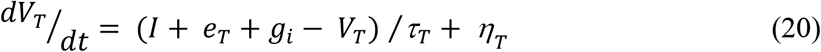

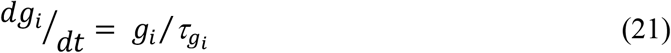

Here, *e* is the equilibrium potential of each unit (*e_i_* = −80 mV, *e_T_* = −60 mV), *τ*is the membrane time constant (*τ_i_* = 7 ms, *τ_T_* = 8 ms), and *η* represents Gaussian independent noise added to each unit with standard deviations of 30 mV. The inhibitory current onto the target neuron is defined by *g_i_*. The inhibitory post synaptic potential was set at −10 mV and driven by spikes in the inhibitory unit through their modeled connectivity with a time constant of *τ_gi_* = 4 ms. Both units fired spikes at a threshold of −30 mV and had refractory periods of 1.5 and 2 ms for the inhibitory and target units respectively.

#### Ground truth datasets

The ground truth datasets were taken from the SpikeForest2 (Magland et al. 2020) website from the Kampff ground truth dataset (Marques-Smith et al. 2018; Neto et al. 2016). The sorting results for MountainSort4, SpyKING-CIRCUS, and KiloSort2 were also downloaded and taken as given from the SpikeForest2 website.

### Spike sorter parameters

The spike sorters evaluated in this paper have many parameters. To simply for our analysis, we used the default parameters of each sorter with the exception of clip width. Clip width is a critical parameter that is highly dependent on the data. KiloSort2 uses a clip width of 2 ms and does not include clip width as a user modifiable parameter. For MountainSort4 and SpyKING-CIRCUS, we used a clip width of 2 ms for our synthetic and cerebellum data. For the same datasets, we used our default FBP parameters (Table 1), as these are generally tuned to the smaller spike waveform widths we observe in our own cerebellar data and used in our simulated datasets. For the ground truth SpikeForest2 Kampff datasets, the default settings for KiloSort2, MountainSort4, and SpyKING-CIRCUS were used as reported by SpikeForest2. We found that the ground truth datasets contained units with much wider spike widths than observed in our cerebellum data and so increased the FBP clip width to [1.0, 2.5] ms for these recordings, but continued to use the remaining default inputs shown in Table 1.

**Table 1.**
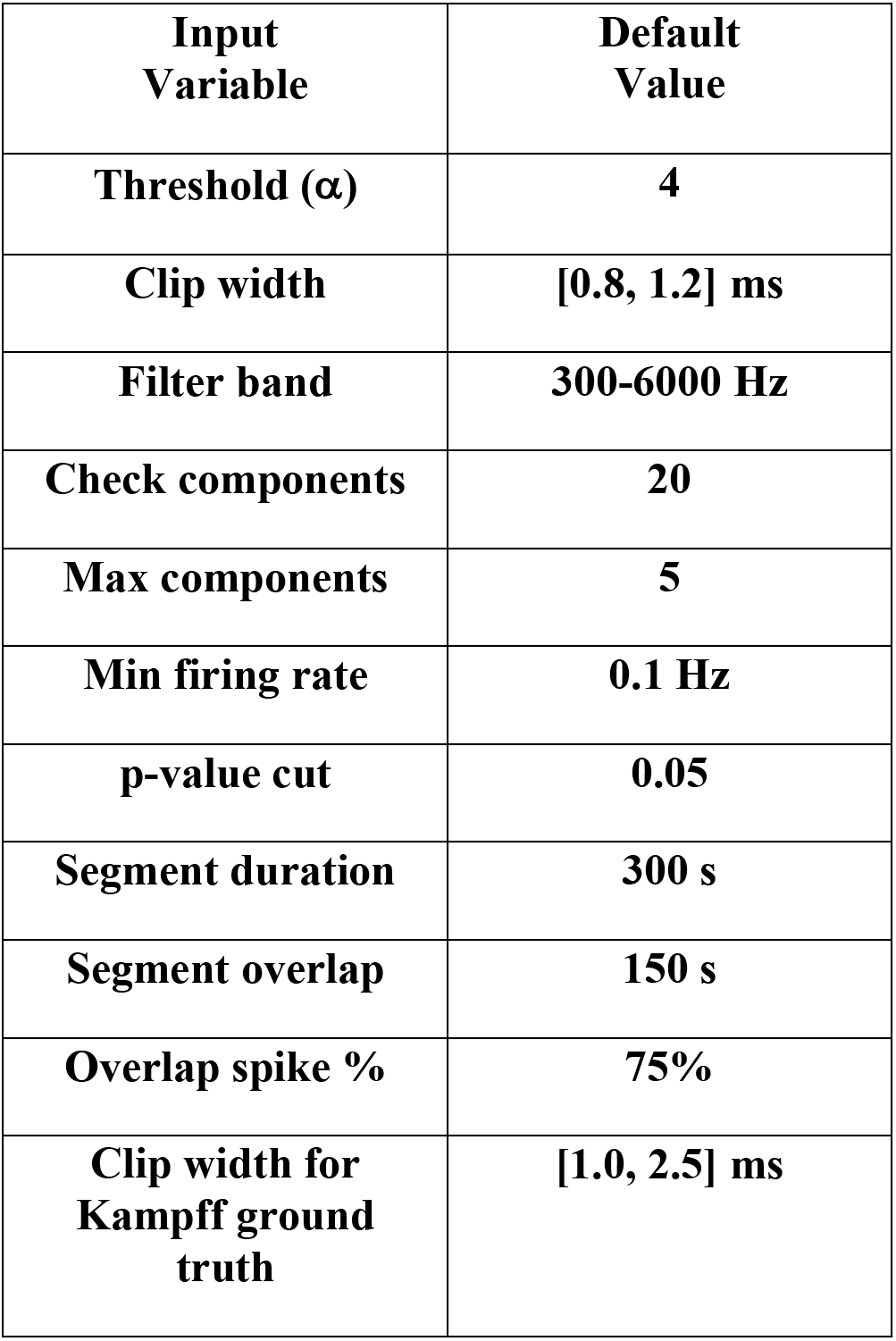
Default parameters of FBP sorter

| Input Variable | Default Value |
| --- | --- |
| Threshold ($\alpha$) | 4 |
| Clip width | [0.8, 1.2] ms |
| Filter band | 300-6000 Hz |
| Check components | 20 |
| Max components | 5 |
| Min firing rate | 0.1 Hz |
| p-value cut | 0.05 |
| Segment duration | 300 s |
| Segment overlap | 150 s |
| Overlap spike % | 75% |
| Clip width for Kampff ground truth | [1.0, 2.5] ms |

### Algorithm implementation

Two key features of our algorithm implementation involve parallelization and could be expanded upon in the future. The clustering procedure runs in parallel processes using the Python Multiprocessing module. Clustering is performed by initiating multiple ‘work items’, each of which sorts a single channel’s data for a single segment. The binary pursuit algorithm runs in massively parallelized fashion on a GPU. Each GPU worker computes the objective function *L*(*t, T*) within a window the width of a clip and stores the results for all windows in GPU memory. In effect, the entire voltage data is chopped up and analyzed in pieces the size of the clip width. Then, a second wave of workers adds a spike in window *W* if *max*(*L*(*t, T*)) in *W* is larger than *max*(*L*(*t, T*)) in both adjacent windows *W-1* and *W+1* and *L*(*t, T*) < = 0 in the next earlier window, *W-2*. The logic for this set of requirements is that subtracting the voltage associated with a spike from the residual voltage in *W* cannot affect any time points beyond its neighboring windows, but the voltage residual in the neighboring window *W-1* is technically not resolved until its neighbor *W-2* is resolved. Thus, we enforce the policy that addition of spikes defers to windows to the left. For the ensuing iteration of binary pursuit, we need to update *L*(*t, T*) only in windows where a spike was added and their neighbors. We then recheck whether or not *L*(*t, T*) > 0 in windows where *L*(*t, T*) either was updated or was greater than zero on the previous iteration. Our strategy tremendously speeds up the binary pursuit algorithm as it reaches convergence, but still is practical only using the massive parallelization afforded by GPU processing. We refer the reader to the source code for precise implementation details.

### Hardware

We used a desktop computer running Windows 10 Enterprise N, 64-bit, with two Intel(R) Xeon(R) Gold 5122 processors (3.60 GHz) with 8 cores each and 384 GB RAM. The graphics card used for GPU processing was an NVIDIA GeForce GTX 1660 Ti with 6 GB dedicated memory.

### Software

The software was written and distributed in Python code using Python 3.7 and additional packages from the standard Anaconda distribution 4.8.3. Parallelization was implemented with the Python Multiprocessing module. The GPU code was written with OpenCL and implemented via the PyOpenCL Python wrapper version 2019.1.2. Several algorithms were further optimized with Cython version 0.29.12.

